# Overlap in neural representations of coordinated wrist and finger movements in human motor cortex

**DOI:** 10.64898/2026.03.19.712976

**Authors:** A.M.X. Emonds, E.V. Okorokova, G.H. Blumenthal, J.L. Collinger, S.J. Bensmaia, L.E. Miller, J.E. Downey, A.R. Sobinov

## Abstract

Dexterous hand function underlies many essential human activities, from tool use to expression through gestures. Coordinated digit movements are enabled by the intricate musculature of the hand and forearm, which also imposes mechanical coupling between the digits and wrist, constraining their independent control. It remains unclear whether motor cortex inherits these constraints in its activity or encodes digit and wrist independently. To address this problem, we asked individuals with intracortical microelectrode arrays implanted in motor cortex to attempt flexion and extension of individual digits, either in isolation or in combination with attempted wrist movements. We could accurately decode which digit was moving based on cortical recordings, and channels selective for digit identity were arranged somatotopically across the recording arrays. Nevertheless, the activity during flexion or extension overlapped between digits, and movement direction of a given digit could be reliably inferred by a decoder trained on movements of other digits. This directional signal was largely invariant to the digit’s initial posture. The population axis describing digit movement direction was aligned with the axes associated with wrist flexion-extension or pronation-supination. This alignment persisted during simultaneous wrist and digit movements, which complicated efforts to control them individually. However, by decoding wrist and digit motion from activity orthogonal to the shared direction axis, a participant was able to achieve continuous control of virtual hand movements with improved speed and reduced unintended movements. Together, the results identify both a code for digit identity and a low-dimensional flexion-extension signal which is shared across the digits and wrist. This arrangement is consistent with muscle-like biomechanical constraints on motor cortical activity, which must be accounted for to improve coordinated BCI control.

## Introduction

The hand endows us with an ability to interact with the world around us, from communicating through gestures to grasping a cup of coffee or playing music. This versatility is enabled by the intricate musculature of the hand and forearm and the disproportionately large regions of sensorimotor cortex devoted to hand control (Penfield and Rasmussen, 1950; Rizzolatti and Luppino, 2001; Baldwin *et al*., 2018; Sobinov and Bensmaia, 2021). Yet, the musculoskeletal architecture that makes dexterity possible also imposes strong biomechanical constraints: the digits share major muscle groups whose bodies are located in the forearm, and tendon connections can transmit force across fingers (Kilbreath and Gandevia, 1994; Reilly and Schieber, 2003; Schieber and Santello, 2004; Duinen and Gandevia, 2011). Moreover, many muscles that actuate the digits cross the wrist, and changes in wrist posture alter the muscle lengths and patterns required to generate digit forces and motion (Adewuyi, Hargrove and Kuiken, 2017; Beringer *et al*., 2020). Predictably, digit-actuating muscles are also active during wrist movements. These mechanics limit the independent control of individual degrees of freedom (Kilbreath and Gandevia, 1994; Zatsiorsky, Li and Latash, 2000; Reilly and Schieber, 2003; Schieber and Santello, 2004; Duinen and Gandevia, 2011). Despite this evidence, an implicit assumption in brain-computer interfaces treats each joint independently, rather than taking interactions into account (Aggarwal *et al*., 2013; Vaskov *et al*., 2018; Shah *et al*., 2023; Jude *et al*., 2026). It remains unclear to what extent motor cortex reflects these mechanics or encodes more independent control signals in BCIs.

A large body of work has shown that motor cortical activity contains information about fine hand kinematics (Schieber, 2001; Capaday *et al*., 2013; Goodman *et al*., 2019; Okorokova *et al*., 2020). Finger and wrist movements activate often overlapping regions within motor cortex (Aggarwal *et al*., 2008; Branco *et al*., 2019; Chehade and Gharbawie, 2023), such that individual neurons within motor cortex are typically involved in the movement of multiple digits (Schieber and Hibbard, 1993; Georgopoulos *et al*., 1999; Acharya *et al*., 2007; Aggarwal *et al*., 2008; Kirsch, Rivlis and Schieber, 2014; Branco *et al*., 2019). This overlap is commonly interpreted as evidence for distributed, mixed selectivity rather than segregation of digit-specific neuronal groups. Despite the apparent somatotopic overlap, decoding analyses suggest partial separability between wrist- and digit-related activity (Mohamed and Aharonson, 2021), indicating that wrist and digit signals should also be separable during simultaneous wrist and digit movements (Nason *et al*., 2021; Mender *et al*., 2024). Importantly, apparent interactions between representations could arise not only from fixed mechanical coupling but reflect functional coupling—statistics of hand use arising from the correlated joint kinematics during everyday behavior (Ejaz, Hamada and Diedrichsen, 2015). Whether neural activity is fundamentally organized around independent digit- and wrist-specific commands, or instead around other activity profiles constrained by biomechanics or use, remains unclear—particularly during movements in which wrist and digits are engaged together, as they typically are during natural grasping and manipulation.

This question is not only fundamental for understanding motor control but also has practical consequences for neurotechnology. People with tetraplegia view restoration of arm and hand movement as fundamental to an improved quality of life (Anderson, 2004; Collinger, Boninger, *et al*., 2013). Intracortical brain-computer interfaces (iBCIs) have enabled functional arm control and simple grasping behaviors (Collinger, Wodlinger, *et al.*, 2013; Ajiboye *et al*., 2017; Flesher *et al*., 2021; Rastogi *et al*., 2021), but more complete restoration of hand function will require coordinated control of individual digits together with the wrist. Although high-density electrocorticography (ECoG) and intracortical recordings have enabled accurate decoding of finger movements in both offline and online settings (Kubánek *et al*., 2009; Hotson, McMullen, Matthew S Fifer, *et al*., 2016; Irwin *et al*., 2017; Jorge *et al*., 2020; Nason *et al*., 2021; Guan *et al*., 2023; Nason-Tomaszewski *et al*., 2023; Vargas-Irwin *et al*., 2024; Willsey *et al*., 2025), achieving high-dimensional and fully independent control remains out of our grasp; systems supporting many degrees of freedom are under active development (Wodlinger *et al*., 2015; Hotson, McMullen, Matthew S Fifer, *et al*., 2016; Nason *et al*., 2021; Shah *et al*., 2023; Dekleva and Collinger, 2025). Moreover, adding coordinated wrist control may induce gain adaptation, reducing effective bandwidth or otherwise altering individual movement representations (Rasmussen, Schwartz and Chase, 2017). Together, these limitations raise the possibility that understanding the constraints intrinsic to motor cortical population activity—and not just decoder design—may be crucial for extracting hand control signals for BCI.

Here, we examined whether human motor cortical activity reflects coupling between digits and wrist or natively supports independent control signals. Using intracortical recordings from participants with tetraplegia, we analyzed population activity during attempted digit flexion or extension, performed either in isolation or concurrently with wrist movements. We asked how digit identity and movement direction are encoded and spatially organized across the arrays, quantified overlap between digit- and wrist-related signals, and exploited this structure to build decoders that enabled continuous control of a virtual hand.

## Results

We recorded motor cortical activity using intracortical microelectrode arrays (Neuroport Array, Blackrock Neurotech, UT, USA) implanted in the motor cortex (Figure 1A) of three individuals with tetraplegia due to cervical spinal cord injury who attempted to make finger and wrist movements following audiovisual instructions (Figure 1B). We examined how this cortical activity represents both digit identity (ID) and the direction of digit motion, as well as the interaction of the representation of digit motion with that of wrist motion.

**Figure 1.**
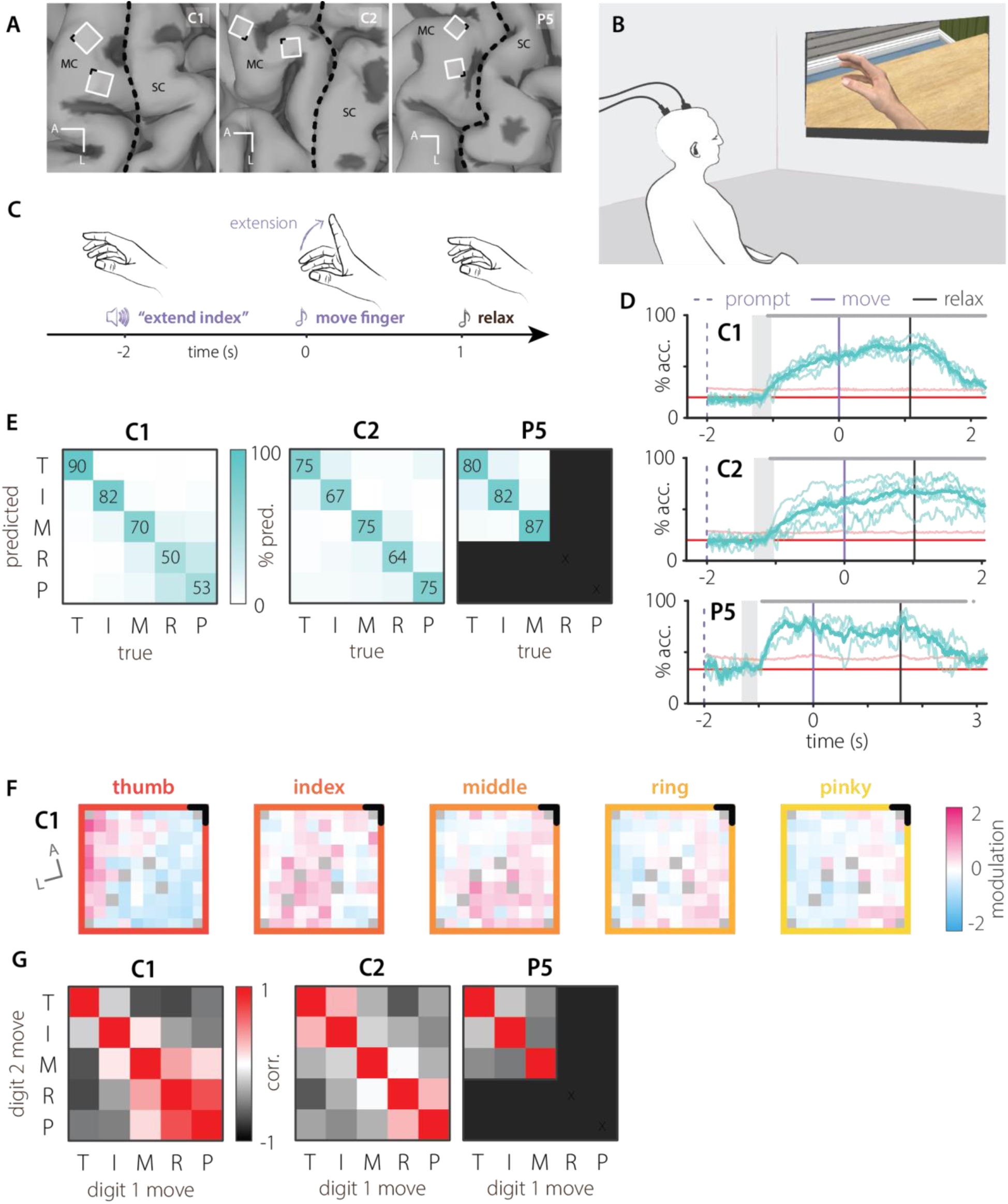
Decoding digit movement. **A|** Implant locations for all participants. **B|** Experimental setup. **C|** Trial progression. **D|** Time course of digit classification accuracy for each participant at each time point for decoders trained on both flexion and extension trials. Each thin trace corresponds to a different session, and thick—to average. Gray bar above the traces corresponds to the points where prediction significantly exceeded chance performance of shuffled distribution (pale pink line). Red line illustrates chance level. Gray shaded region represents the range in which audio cues ended. **E|** Digit identity confusion matrices constructed at the peak accuracy and averaged across sessions. **F|** Modulation of electrodes by digit identity (significance by Kruskal-Wallis test, see **Supp. Figure 1C** and details in Methods) on C1 lateral motor array during single digit movement, averaged across sessions. Pink corresponds to facilitation of activity and blue, to inhibition. Non-significant channels are gray. Anterior-most medial corner of the array is black. (C1 n=5, C2 n=6, P5 n=3) **G|** Similarity of spatial modulation maps for each subject. Since these correlations were calculated using composite maps built across all sessions, the diagonal represents a comparison between identical maps.

### Movement of individual digits

We investigated the neural activity underlying individual finger movements by asking participants to mimic the movements of a virtual hand, which they observed on a monitor (Figure 1B). Trials started with the participants imagining their hand and wrist in a neutral posture. An auditory instruction specifying a random digit and movement direction was followed two seconds later by an auditory “go” cue and the movement of the virtual hand, which the participant was instructed to mimic (Figure 1C). A subsequent “relax” cue instructed them to return to the neutral posture. We built linear decoders on neural activity from each channel, averaged within 100ms windows, only selecting channels with a mean firing rate above 0.5 Hz. Neural activity allowed accurate classification of digit identity starting from 125ms, 125ms, and 265ms after audio cue end (1.08s, 1.08s, and 940ms before the go cue) in C1, C2, and P5, respectively (Figure 1D), with accuracy peaking after the go cue. In participant C1, discriminability of the digits generally decreased from thumb to pinky (Figure 1E), but in other participants the accuracy was similar for all digits, although P5 only imagined thumb, index and middle digit movements. Overall, digit identity was strongly represented in the neural activity of all participants.

We hypothesized that differences in discriminability might reflect the spatial organization of modulated channels on the arrays. If so, digits that were more frequently confused would be expected to activate highly overlapping regions of motor cortex. We made modulation maps by computing the firing rate of a channel relative to its average firing rate during the movement phase of the trial (Figure 1F, Supp. Figure 1A). In all participants, we focused the spatial analysis on the more active array—lateral in C1 and C2, and medial in P5—because the other array was mostly silent (mean firing rate < 0.5 Hz). We selected only the channels with modulation that differed consistently across digits (Kruskal-Wallis test, p<0.05). The positively modulated units were somatotopically organized in all participants, most saliently in C1 (Figure 1F), such that movements of neighboring digits recruited activity in nearby channels more than distant ones. For each participant, we quantified the similarity between the modulation maps for individual digits by computing the spatial correlation between them (Figure 1G). We identified significant correlations by comparing them with correlations obtained from shuffled data pooled across all conditions (Supp. Figure 1B). The patterns of spatial correlation in all participants qualitatively matched the decoding confusion matrices (Figure 1E and Figure 1G, respectively). The confusion in decoding middle, ring, and pinky in C1 (Figure 1E, left) was reflected in high spatial correlation between the maps for those digits (Figure 1G, left). At the same time, high discriminability of the thumb was reflected in low or negative spatial correlations. In C2, confusion between the thumb and index was reflected in the spatial correlations of their associated maps. We did not see these effects in P5, most likely because fewer digits were tested across fewer sessions, lowering the total number of significantly modulated electrodes. Overall, spatial correlations between neighboring digits were slightly higher than between digits that were far apart, suggesting not only spatial organization of neural activity corresponding to each digit, but also a certain degree of somatotopy across digits.

Having characterized the distinct patterns of activity corresponding to digit identity, we next examined how digit movement direction is encoded in the motor cortex. We isolated trials corresponding to the movement of a single digit and built decoders to indicate whether that digit was flexed or extended (Figure 2A). In participant C2, movement direction was represented much more weakly than in the other two participants, and signals were consistently weak for all digits (**Error! Reference source not found.**Supp. Figure 2A). In the other two participants, C1 and P5, movement direction information appeared at a similar time asdigit identity (1.12s and 820ms before movement, respectively). The encoding of movement direction did not strongly depend on the starting posture of the movement. Decoders trained on the “move” phase—where flexion and extension started from neutral posture—largely generalized to the “relax” phase, where movement started at one of the extremes of the range of motion, and vice versa (same starting posture mean accuracy 91%, different starting posture mean 84%, p=0.0026, Figure 2B; see Supp. Figure 3 for temporal generalization heatmaps).

**Figure 2.**
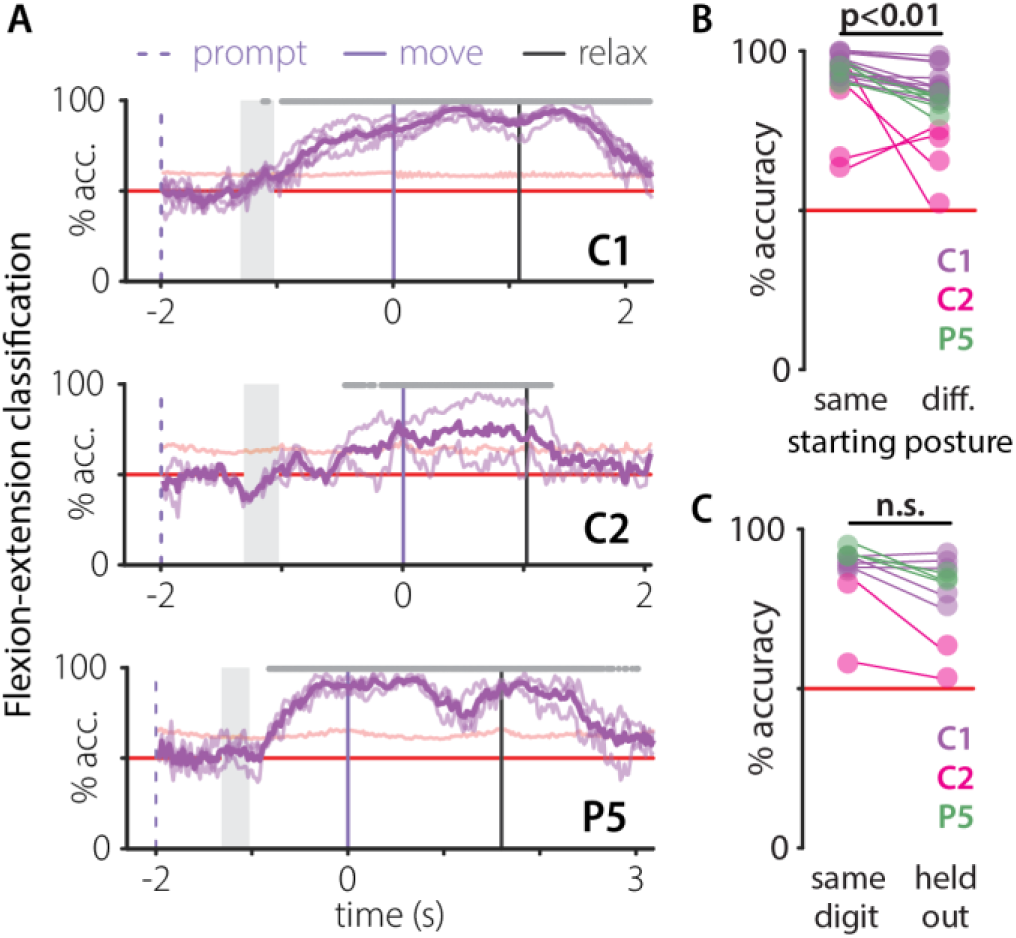
Movement direction generalizes between digits. **A|** Average classification of digit movement direction—flexion or extension—for individual digits. Lines and shading follow the conventions defined in Figure 1D. (C1 n=5, C2 n=2, P5 n=3) **B|** Generalization of digit movement direction encoding for different starting postures. Decoders were trained on the 100ms window of activity during movement that maximized classification accuracy. Different colors correspond to different subjects (C1: purple, C2: pink, P5: green). Thin lines connect performance during the same session. **C|** Generalization of movement direction across digits during movement phase in subject C1. The extension-flexion classifier was trained and tested on single digits (“same digit”) and trained on all but the tested digit (“held out”). Each dot represents average accuracy in a single session. Colors as in B. Thin lines connect sessions. Horizontal solid red line corresponds to chance level.

Given the biomechanical coupling between digits, we assessed whether movement direction of one digit could be decoded from the neural activity corresponding to movement of other digits. We built decoders of movement direction on trials with all but one digit and compared them to decoders built on only that digit (Figure 2C; for elaboration, see Supp. Figure 4). The decoders tested on the untrained digits performed better than chance, and not significantly worse than the same-digit decoders (p=0.058). Flexion or extension of each digit evoked activity in channels widely spread over the arrays of the participants (Supp. Figure 2BC). Overall, these results indicate strong digit movement direction encoding in motor cortical activity that generalizes between postures of the digit, as well as across digits.

### Neural overlap between wrist and digits

Digit movement rarely happens in isolation from wrist movements, which are typically needed to optimize hand orientation, as when the hand approaches an object (Yan, Sobinov and Bensmaia, 2022). Also, the powerful extrinsic finger muscles that cross the wrist create substantial wrist torque that must be counteracted by wrist extensor muscles. Furthermore, altered wrist posture changes the force production capacity of the extrinsic hand muscles. The extent to which these complex mechanics are represented within neural activity is an important basic neuroscientific question with significant implications for iBCI control.

We began by characterizing the neural activity during isolated wrist movements. Participants attempted to flex, extend, pronate, or supinate their hand starting from the neutral posture (Figure 3A). Linear classifiers based on neural activity could distinguish wrist orientation throughout these trials, beginning 73ms, 413ms, and 373ms after mean audio cue end (1.20s, 880ms, and 920ms before the movement cue) for C1, C2, and P5, respectively (Figure 3B). Classification accuracy was generally lower in C2 than in C1 and P5, but no consistent confusion pattern appeared across participants (Figure 3C).

**Figure 3.**
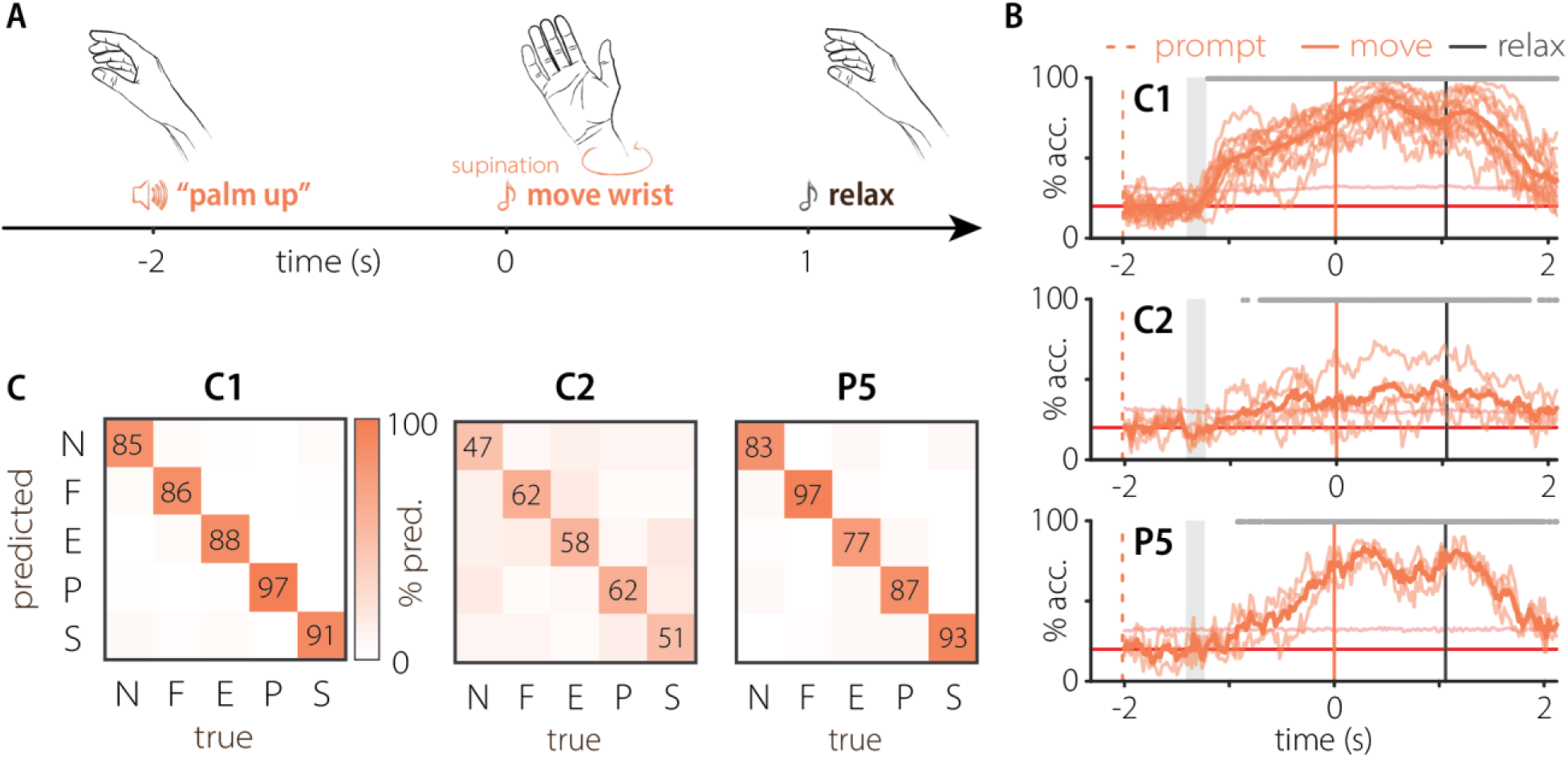
Wrist movements. **A|** Schematic of a representative trial. **B|** Time course of wrist orientation classification accuracy for subject C1 (n=13), C2 (n=4), and P5 (n=3) based on the neural activity averaged in 100ms window at each time point. Lines and shading follow the conventions defined in Figure 1D. **C|** Wrist posture confusion matrix at peak accuracy from B, averaged across sessions.

We then looked at the interaction between neural activity for isolated wrist and digit movements by comparing movement axes in neural space. This analysis was performed with data from C1 and P5, who had strong representation of digit movement direction (Figure 2, Supp. Figure 3C) and wrist movement direction (Supp. Figure 5) throughout the trials. We defined a digit movement direction axis in neural activity space as the vector running from neural activity during extension through the neutral posture to activity during flexion. We identified digit movement direction axes both for each digit individually and for all digits combined, and computed angles between the axes for each digit and the common axis (Figure 4A). In C1, the vectors for individual digits aligned closely with the common digit movement axis, which was expected given that digit movement direction generalized between digits (Figure 2C, purple dots). Predictably, this was less pronounced in P5, whose digit movement direction generalization, while significant, was weaker than that of the other participants (Figure 2C, green dots). This was possibly due to a learned compensatory strategy P5 had developed that relied upon residual movement ability in the index finger and thumb.

**Figure 4.**
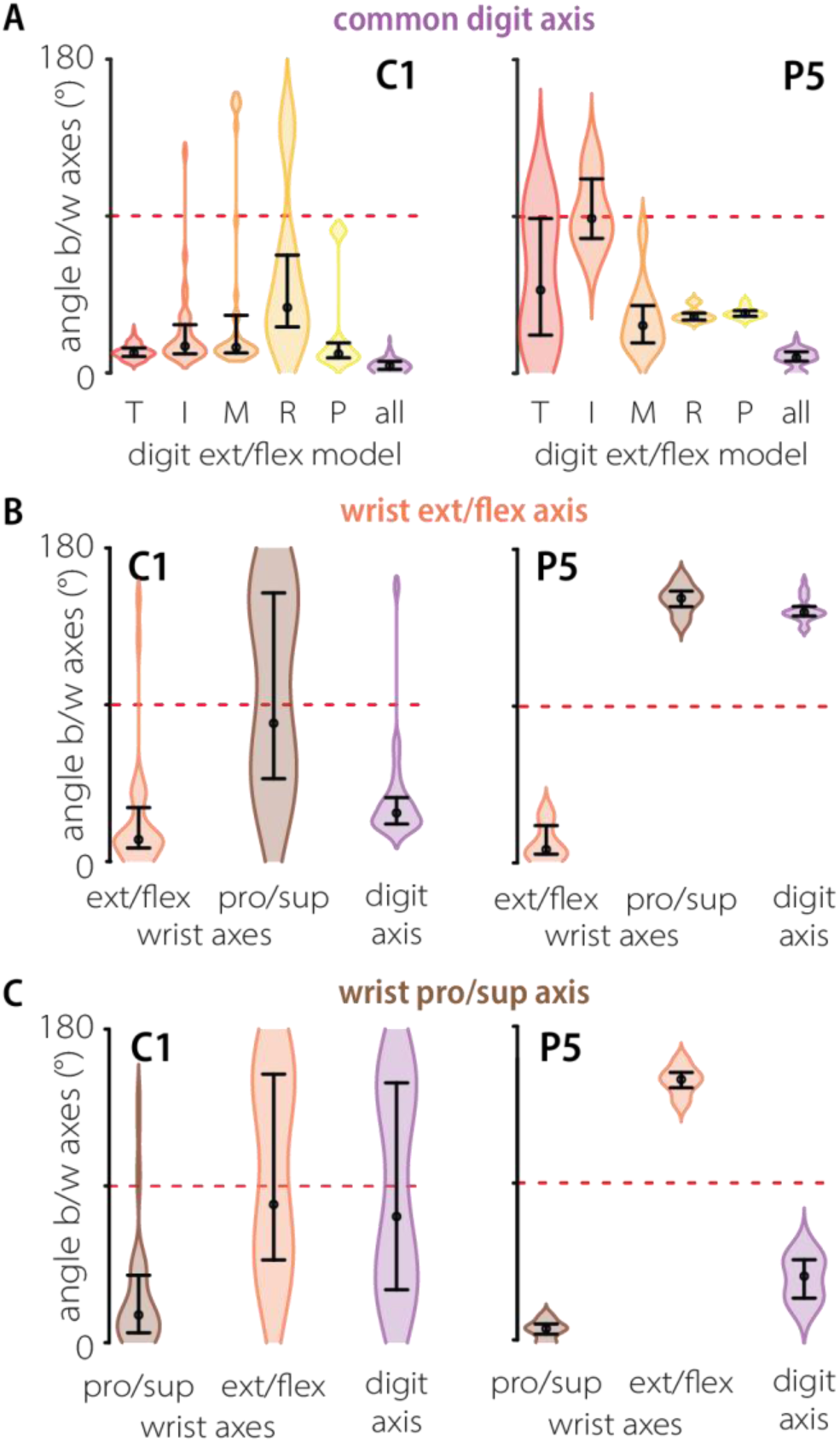
Overlap between digit and wrist neural axes. **A|** Distribution of angles between a common neural digit movement direction axis and the same axis computed for each digit independently (C1 n=6 and P5 n=2 for T, I, M, and all; C1 n=1 and P5 n=1 for R, P). “all” corresponds to angles between models computed on different data folds. Black dots represent the median and bars depict the interquartile range (IQR). A dotted red line at 90° denotes the vector angle of orthogonality. T: thumb; I: index; M: middle; R: ring; P: pinky. **B|** Distribution of angles between wrist extension-flexion (EF) axis and wrist supination-pronation (SP) axis and common digit EF axis. (C1 ext/flex n=5, pro/sup n=6; P5 n=2) **C|** Same as B for the wrist supination-pronation axis.

Similarly, we defined neural axes for wrist extension to flexion (EF) and supination to pronation (SP), and calculated angles between those axes and the common digit axis from data collected on the same day. Wrist EF and SP neural axes were orthogonal in participant C1 (Figure 4B, brown and orange distributions), which was expected since they correspond to orthogonal directions of movement and are largely actuated by different muscles. Wrist EF and digit EF axes were aligned (Figure 4B, purple), but wrist SP and digit EF were orthogonal (Figure 4C, purple). One possible explanation is that finger movements predominantly evoke flexion and extension torques at the wrist, with little contribution to pronation or supination. In participant P5, the EF and SP neural axes were aligned, but pointed in opposite directions. Although the sources of these interactions in the neural space are unclear, the neural activity underlying movements of the wrist and digits is clearly linked.

### Simultaneous movement of wrist and digits

Given the interaction between wrist and digit encoding during independent movements, we asked whether the motor cortex would separate them when participants attempted to move the wrist and digits simultaneously. After two sequential instructional cues—first wrist and then digits—a single tone cued participants to attempt movement of both wrist and a specified finger at the same time (Figure 5A). The different wrist orientations, digit identities, and digit movement directions were randomized and interleaved. While the timing of digit-identity information was similar to digit-only trials (1.00s and 960ms before movement and 205ms and 245ms after mean audio cue end for C1 and C2, respectively, Figure 5B) and peak digit identity classification accuracies were similar (Supp. Figure 6A, p=0.41), classification of digit identity was poorer during the preparatory period during simultaneous movements (Supp. Figure 6B, p<0.0003). A similar pattern was observed for the wrist: similar latency (2.24s and 1.60s before movement, 33ms and 673ms after mean audio cue end for C1, C2, Figure 5C) but reduced overall accuracy (Supp. Figure 6C, p=0.0023), particularly during preparation (Supp. Figure 6D, p=0.0038).

**Figure 5.**
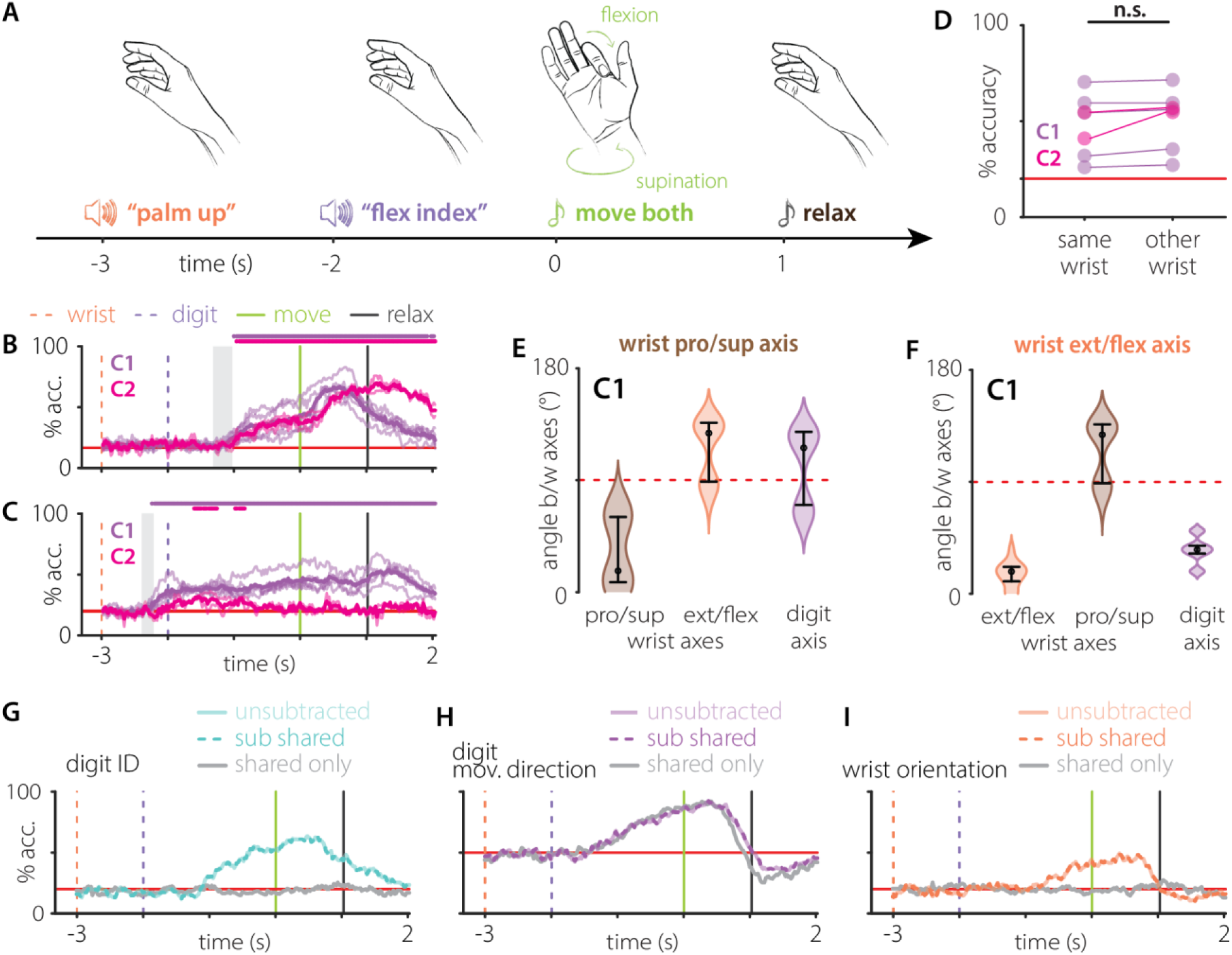
Simultaneous wrist and finger movements. **A|** Schematic of an example trial. **B|** Time course of digit identity classification accuracy for subjects C1 (purple) and C2 (pink) based on the neural activity at each time point. Lines and shading follow the conventions defined in Figure 1D. **C|** B, for wrist orientation classification. **D|** Mean digit classification accuracies within the same wrist orientation (“same wrist”) and across wrist orientations (“other wrist”) for each participant (colors as in B and C). Wilcoxon rank sum test p>0.05 within each session and with all sessions pooled. Breakdown by session in Supp. Figure 6E. Breakdown by train and test wrist orientation pair in Supp. Figure 6F. (C1 n=5, C2 n=2) **E|** Distribution of angles between wrist supination-pronation axis, wrist extension-flexion axis, and common digit extension-flexion axis (C1 n=1). **F|** E, for the wrist extension-flexion axis. **G|** Classification accuracy of digit identity by a decoder trained on movement using full neural space (pale teal), shared axis-subtracted neural activity (dotted teal), and neural activity along the shared axis alone (gray). Train and test trials are matched across decoders. **H|** F, for digit movement direction. **I|** F, for wrist orientation.

To further probe finger and wrist interactions, we evaluated whether wrist orientation affected digit identity signals in the motor cortex. We trained decoders of digit identity on one wrist orientation and tested them on other orientations (Figure 5D). Surprisingly, wrist orientation had no effect on the digit decoding accuracy in either participant (p=0.45, breakdown by session and training/testing wrist orientation in Supp. Figure 6EF)

We then investigated whether the combined movements of the digits and wrist affected the previously discovered neural geometry. For that, we computed movement direction axes corresponding to neural spaces for digit extension to flexion (digit EF), wrist extension to flexion (wrist EF), and wrist supination to pronation (wrist SP) and calculated angles between those axes. Consistent with our expectations from previous experiments, the SP axis was fully orthogonal to the digit and wrist EF axes (Figure 5E). The wrist EF axis remained aligned with the digit EF axis (Figure 5F), even for the dataset containing all possible combinations of wrist and digit movements, underscoring the anatomical and functional connection between wrist and digit flexion-extension movements.

However, this large overlap could impact our ability to decode desired movements during BCI control and lead to unintended movements. It would also prevent us from training decoders on separate sets of digit and wrist movements, forcing exploration of all possible combinations. As a solution, we removed the shared axis between wrist and digits from neural activity and extracted task information from the remaining neural space. Specifically, we removed the component of activity aligned with the common digit axis. Even in this new, reduced neural space, we could decode which digit was moved and its direction of motion (Figure 5GH). Furthermore, the space retained reliable information about intended wrist movement (Figure 5I). Restricting the analysis to this neural subspace allowed for reliable decoding of movement intent and could be completed without the need to sample all possible combinations of wrist and digit movements.

### Online decoding

Building on the principles learned in offline analyses, we implemented an online paradigm to allow participant C1 to control a virtual robotic hand. We started by training Kalman filter-based position and velocity decoders as a participant observed 10-20 repetitions of thumb, index, or middle finger flexion or extension. The participant was then asked to move the digit to the instructed target posture within 15 seconds while maintaining the other digits in the neutral position, before returning all digits to the neutral posture (Figure 6A and Supp. Figure 7A, green and gray regions). Testing first with decoders trained on the full neural space, C1 achieved a success rate above 80% (move success 94% (281/300), return success 81% (218/268), Supp. Figure 7AB). However, movement times were long (median times of 2.5s, IQR 1.8-4.7s to move to the target and 4.4s, IQR 2.9-7.2s to return, Supp. Figure 7A). We next built decoders on the neural space without the common movement direction axis. Under this condition, C1 could achieve target postures and return to neutral significantly more quickly (to the target: median 2.2s, IQR 1.5-4.2s, p=0.0073, Figure 6B purple swarms; to return to neutral: 3.5s, IQR 2.8-5.6s, p=0.0122, Figure 6B black swarms; full traces in Figure 6A). Furthermore, it improved the success rate of movements to the target to 95% (199/210) and of returns to 96% (191/198) (Figure 6C).

**Figure 6.**
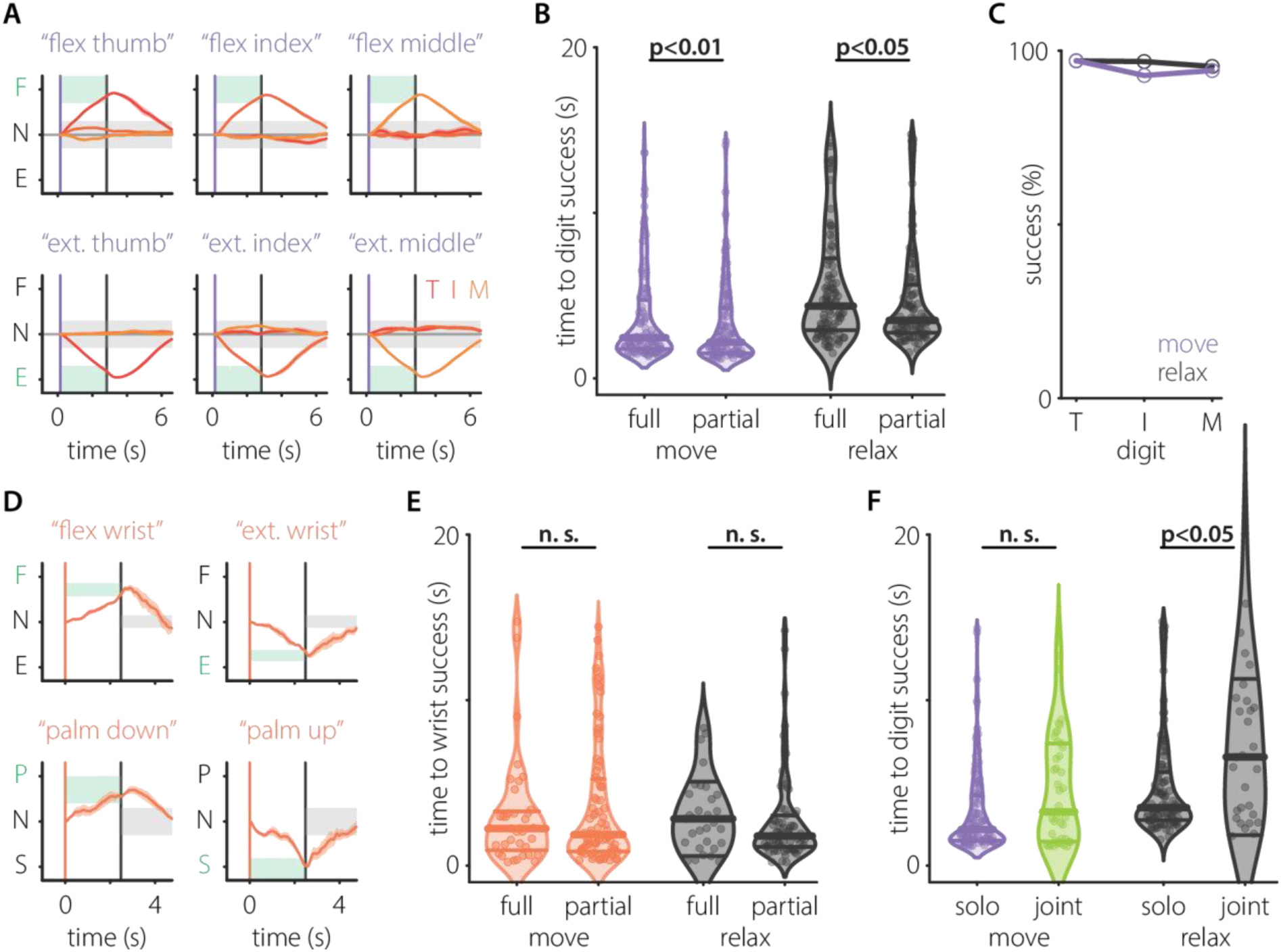
Online control. **A|** Position during successful individual finger movement trials completed using a shared axis-subtracted decoder, averaged by target (C1 n=3) and time-warped to the median stage length across successful trials. Target window is shaded green. Gray regions represent error threshold for non-target digits and for target digit during return. Red and orange shaded regions surrounding traces depict the standard error of the mean. **B|** Distributions of time to digit “move” (purple swarms) or “relax” (black swarms) phase success for C1 using a decoder built on the full neural space (“full”) versus a decoder built on the common axis-subtracted neural subspace (“partial”). Median represented by thicker and IQR represented by thinner horizontal lines. **C|** Trial success rates during movement (purple trace) and relaxation (gray trace) phases for individual finger movements in A, broken down by digit. **D|** Extension/flexion (top row) and pronation/supination (bottom row) during successful wrist movement trials completed using a shared axis-subtracted decoder, averaged by target (C1 n=2). Interpolation and target window shading as in A. Orange regions surrounding traces depict the standard error of the mean. **E|** B, for wrist experiments. Movement phase swarms now depicted in orange. **F|** Distributions of time to digit “move” or “relax” phase success for C1 using a common axis-subtracted decoder while performing individual finger movements (“solo,” purple/black swarms) versus simultaneous wrist and finger movements (“joint,” green/black swarms).

To investigate wrist movement control, we asked the participant to extend, flex, pronate or supinate their hand, then return to neutral posture. We trained Kalman filter-based position and velocity decoders on common-axis-subtracted neural space to determine whether the orthogonal subspace contained enough information for wrist control. On average (median), C1 achieved the target postures within 1.9s (IQR 0.9-5.2s) and returned to neutral within 1.8s (IQR 1.1-3.0s, Figure 6D). This was on par with performance in full neural space (p=0.73 and p=0.15 Figure 6E, orange and black swarms, respectively; Supp. Figure 7CD). C1 achieved an 86% (85/99) success rate during movement and 93% (74/80) during return (Supp. Figure 7E).

Lastly, we explored the ability of the participant to make concurrent wrist and digit movements by training digit movement decoders on digit-only trials and a wrist decoder on wrist-only trials after subtracting the common extension-to-flexion axis. The participant was asked to flex or extend a single digit and orient the wrist at the same time. They successfully completed 83% (45/54) of movement phases and 69% (29/42) of returns, indicating that C1 can selectively flex and extend digits during wrist movements (Supp. Figure 7FG). Although we could not directly enforce the concurrency of the movements beyond instructing the participant, movement traces show that all joints moved toward the target posture together (Supp. Figure 7H). Due to weak wrist decoder performance on the day of testing and unbeknownst to the participant, however, the decoder received assistance from the computer with wrist orientation. As in the separate control experiments, C1 moved the target finger relatively quickly (1.5s, IQR 1.2-2.6s during movement and 4.1s, IQR 3.0-6.6s during relaxation), but stabilizing the other digits and wrist took an additional 1.8 and 2.5 seconds (Supp. Figure 7H, see Supp. Figure 7I for corresponding wrist traces). By subtracting the common axis from the full neural population activity, we derived decoders that could be trained independently on each movement group. This approach also reduced unintended digit and wrist movements, bringing the time to success to the level of separate digit movements (p=0.099, Figure 6F purple and green swarms), reduced the dependence of finger-movement representations on concurrent wrist movement, and ultimately enabled C1 to control wrist and digit movements simultaneously.

## Discussion

Dexterous hand function relies on neural control signals operating a musculoskeletal system that couples the digits and wrist through shared tendons and multiarticular muscles. Here we show that there is a structured overlap across representations of digit and wrist movements in motor cortex as well, challenging an often implicit assumption that joints are represented independently. These shared subspaces contained the direction of movement of each joint, consistent with anatomical organization of wrist and digit musculature.

Movements of individual digits are rarely fully independent: attempting to move or produce force with one finger typically evokes unintentional motion and/or force in other digits (Kilbreath and Gandevia, 1994; Hager-Ross and Schieber, 2000; Zatsiorsky, Li and Latash, 2000; Lang and Schieber, 2004). Peripherally, this interdependence is explained at least in part by shared musculotendinous anatomy. Multi-tendoned extrinsic hand muscles—such as *flexor digitorum profundus* and *flexor digitorum superficialis*—send tendons to multiple digits, with incomplete functional subdivision across fingers. Connective tissue links can also transmit forces between digits (Reilly and Schieber, 2003; Schieber and Santello, 2004; Duinen and Gandevia, 2011; Lemelin and Diogo, 2016). Coupling is also evident within the spinal neural circuits that contribute to hand actions: spinal interneurons contribute to activation of multiple hand muscles and are thought to support coordinated multi-muscle activation patterns during precision grip (Takei and Seki, 2010; Takei *et al*., 2017). One might expect that, upstream, the movement of separate joints would be abstracted and independent from each other. However, in our data from motor cortex, we observed strong interactions between neural signals associated with digit flexion and extension, with movement direction generalizing across digits. This instead supports the view that motor cortical population activity embeds some properties of the biomechanical constraints on digit control. We further found that the direction of digit movement (flexion versus extension) was agnostic to the digit’s starting postures. This is notable given that, during grasping, hand-related motor cortical activity carries prominent posture-related signals (Goodman *et al*., 2019; Agudelo-Toro *et al*., 2024), which are more informative than velocity. In contrast, reaching-related activity in M1 reflects arm movement velocity (Moran and Schwartz, 1999; Paninski *et al*., 2004; Wang *et al*., 2007; Weber *et al*., 2011). However, even when posture-related modulation is strong, it does not fully obscure directional information. Ultimately, this posture-tolerant directional signal is consistent with the peripheral organization of digit control: flexion and extension of individual fingers are driven predominantly by the same flexor and extensor muscles, even though their mechanics vary with posture. Appropriately, our Kalman filter-based linear decoder performs best when trained on (and predicting) both joint angle velocities and positions.

We observed a somatotopic organization of digit identity across the recording arrays, which may seem surprising given extensive evidence that digit representations in human motor cortex overlap substantially (Donoghue, Leibovic and Sanes, 1992; Schieber and Hibbard, 1993; Dechent and Frahm, 2003; Rathelot and Strick, 2006). Importantly, strong overlap does not preclude somatotopy (Hluštík *et al*., 2001; Indovina and Sanes, 2001). While we also observed broad overlap across digit-related activity, by subtracting the average activation signal, we revealed digit-specific hotspots of activity. These findings extend prior work demonstrating spatial organization of digit-related signals in human motor cortex, including ECoG and high-density surface recordings, and show that penetrating recordings can resolve digit-specific spatial organization in humans (Miller *et al*., 2009; Wang *et al*., 2009; Hotson, McMullen, Matthew S. Fifer, *et al*., 2016; Shelchkova *et al*., 2023).

Digit movements are often accompanied by the movement of the wrist (Yan, Sobinov and Bensmaia, 2022). Complementing digit decoding, we could also decode intended wrist movements from activity in the motor cortex. The strength of this representation varied between participants, however, consistent with sensitivity to recording site and the specific neural populations sampled by each implant (Vargas-Irwin *et al*., 2010; Ajiboye *et al*., 2012). The wrist flexion-extension axis showed strong alignment with the shared digit flexion-extension (EF) axis, and this alignment was consistently greater than that between digit EF and wrist supination-pronation (SP). This is consistent with peripheral biomechanics: the largest extensors and flexors of the digits cross the wrist, generating a torque that needs to be counterbalanced by the wrist muscles. Similarly, changing the wrist posture affects the muscle activity producing digit movements and grasping (Seo *et al*., 2008; Beringer *et al*., 2020; Popp *et al*., 2023).

These effects differ between wrist movements—supination-pronation of the wrist is produced by muscles that do not cross the wrist and should interact less with the musculature that actuates the digits. However, we did observe interaction between the associated activity in one of the participants. Unexpectedly, the digit EF axis aligned with wrist EF in the same direction rather than the opposite direction, as might be anticipated from purely compensatory wrist torques during finger flexion. A possible explanation is that the shared population dimension that we read from the neural activity largely reflects the digit flexion-extension muscle drive. These would flex during wrist flexion and extend during wrist extension and thus naturally covary with wrist flexion-extension, producing the observed alignment in the neural space.

Although we did not see an effect of wrist orientation on digit identity decoding, we consistently observed reduced decoding accuracy of the digits and wrist in preparation for and execution of simultaneous movements. One explanation is limited bandwidth in the sampled motor cortical population: as task complexity increases, the depth of modulation related to particular features within the task may be reduced, an effect seen in monkeys making 2D and 3D virtual reaches (Rasmussen, Schwartz and Chase, 2017; Ferguson and Cardin, 2020). This effect parallels normalization computations widely observed in sensory systems, where responses to a feature can depend on overall stimulus strength, e.g., brightness or volume (Carandini and Heeger, 2012).

Finally, by accounting for the shared structure of digit-related neural activity, we built decoders that improved the ability of our participant to control multiple digits at the same time: flexing or extending one digit while not moving the others. This result highlights how the overlap in neural spaces can be managed rather than treated purely as noise. This performance is encouraging, but it does not yet achieve the precision, stability, or dimensionality of truly dexterous hand control. Our results complement and extend other efforts in restoring hand and arm control (Collinger, Wodlinger, *et al*., 2013; Ajiboye *et al*., 2017; Nason *et al*., 2021; Willsey *et al*., 2025) by focusing on the neural activity underlying simultaneous control of many digits and the wrist. Given the constraints of the recorded signal induced by the spatial organization of informative channels, achieving fully dexterous control will likely require larger spatial sampling, including caudal M1 hidden in the central sulcus, which contains circuitry more directly linked to distal musculature thought to be critical for individual finger movement (Lemon and Griffiths, 2005; Rathelot and Strick, 2009). It may also benefit from recordings across multiple cortical areas that carry complementary information about hand movements (Guan *et al*., 2023).

Here, using intracortical microelectrode arrays implanted in the motor cortex of human participants, we characterized the representation of individuated digit and wrist movements both during isolated and simultaneous movement attempts. We found somatotopic organization of digit identity across the arrays alongside robust digit- and wrist-related signals. We also identified a shared low-dimensional axis in the motor cortex describing digit and wrist flexion-extension that can confound independent decoding. Exploiting orthogonal dimensions enabled better digit BCI performance, including during simultaneous wrist movements. This decoding approach should ultimately enable skilled dexterous control of a prosthetic hand for people with upper limb motor impairments.

## Methods

### Participants

Participants gave informed consent prior to participation in a clinical trial (NCT01894802) conducted under an investigational device exemption from the US Food and Drug Administration and with approval from the Investigational Review Boards at the Universities of Chicago and Pittsburgh. Participants C1 and C2 had C4-level ASIA D spinal cord injuries resulting in paralysis of their dominant right hand. Participant P5 had C8-level ASIA A spinal cord injury resulting in a paralysis of the dominant right hand. All three participants had two 96-channel microelectrode arrays (Blackrock Neurotech, Salt Lake City, Utah) implanted in the left motor cortex. Participant C1 had arrays implanted 35 years after injury occurred and 41-57 months before the experiments reported here. Participant C2 had arrays implanted 4 years after injury occurred and 11-27 months before the experiments reported here. Participant P5 had arrays implanted 6 years after the injury and 3-5 months before the experiments reported here.

### Classification and significance testing

To identify the presence of the information about the condition (e.g., which digit was being moved, or wrist posture assumed), we have trained linear discriminant analysis (LDA) models on each session separately. We have used five-fold stratified cross validation to avoid overfitting. The thin lines in plots like Figure 1D represent classification accuracy traces across a single session. The mean accuracy across sessions is illustrated with a thicker line.

To determine significance, a null distribution of raw classification accuracies (*null distribution_t_*) was created at each time point by training 100 models per session on trials with randomized target assignments and testing on held-out trials from the same session (80/20 split with five folds of cross-validation). The mean classification accuracy trace across sessions was computed, and significance at each time point was determined as

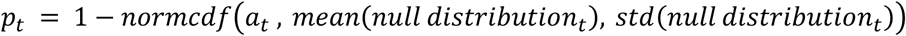

where 𝑎_𝑡_ represents the mean classification accuracy at time 𝑡 and 𝑝_𝑡_ represents the p-value at time 𝑡. To compensate for multiple comparisons, classification accuracy was considered to be significantly above chance where the Benjamini & Hochberg-adjusted p<0.05 (pink horizontal line in e.g. Figure 1D). Specific stage durations varied slightly across sites, as is evident in the stage timing differences between C1/C2 and P5 in Figure 1D and Figure 2A.

### Measuring appearance of conditional information relative to mean prompt completion time

The instruction cue marker in all experiments represents the time at which the audio prompt began to play, not the time at which the entirety of the audio prompt had been delivered. As such, the appearance of information about the task conditions in all cases was occasionally reported relative to the mean prompt completion time for each experiment—the time of the instruction cue plus the mean duration of the audible portion of the prompt as determined from the relevant .wav files.

### Classification generalization matrices

To produce time generalization matrices like the one depicted in Supp. Figure 3C, a series of linear classifiers were trained on a sliding window of 100ms of neural activity beginning at time x and used to predict the task condition, e.g., digit movement direction, for each time point over the entirety of the time course (y). Training and testing data were 80/20 split across trials with five folds of cross-validation. Supp. Figure 3C and Supp. Figure 5A-D represent mean heatmaps of temporal generalization across sessions.

To emphasize the negative deviation from chance signifying decoded movement in the opposite direction in two-class models, Supp. Figure 3C and Supp. Figure 5A-D are presented in units of adjusted accuracy, where 𝑎𝑑𝑗𝑢𝑠𝑡𝑒𝑑 𝑎𝑐𝑐𝑢𝑟𝑎𝑐𝑦 = 2 ⋅ (𝑟𝑎𝑤 𝑎𝑐𝑐𝑢𝑟𝑎𝑐𝑦 − 0.5). An adjusted accuracy of 0 means that classification is at chance level. 1 means classification is always consistent with trial type. -1 means classification is always opposite the trial type.

To determine the threshold of significance conveyed in each heatmap colorbar, 100 linear classifiers with randomized targets were trained on a sliding window of 100ms of neural activity beginning at time x and used to predict e.g. digit movement direction over the entirety of the time course. Two-tailed significance at each time point in a heatmap of adjusted accuracies was determined as

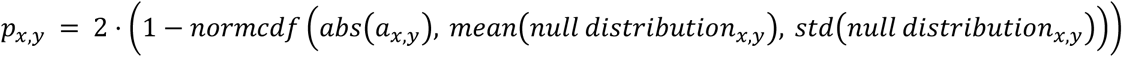

where 𝑎_𝑥,𝑦_ represents the adjusted accuracy at train time 𝑥 and test time 𝑦 and 𝑝_𝑥,𝑦_ represents the p-value at train time 𝑥 and test time 𝑦. As before, the threshold of significance was taken as the minimum across all x and y of the values of 𝑎_𝑥,𝑦_ producing a Benjamini & Hochberg-adjusted p<0.05.

### Digit and wrist classification axes

To identify the axes in the neural space defining the extension to flexion movement of digits, we first defined the periods corresponding to neutral, flexion, and extension of a digit. For neutral, we used the first 25% of the presentation phase (before completion of the audible instruction). For flexion, we used the first 50% of the movement phase in flexion trials, and the first 50% of the relaxation phase in extension trials. Similarly, for extension, we used the first 50% of the movement in extension trials, and 50% of relaxation in flexion trials. Flexion during movement or relaxation was assigned a kinematic value of 1, extension was assigned a kinematic value of -1, and time points pre-instruction were assigned a kinematic value of 0. Then, we trained an LDA model on those periods, extracting the neural axis from it.

Models derived from the Individual Finger Movement data (Figure 2) were trained on (1) thumb extension-flexion trials only, (2) index extension-flexion trials only, (3) middle extension-flexion trials only, (4) ring extension-flexion trials only, (5) pinky extension-flexion trials only, or (6) all-digit extension-flexion. Models derived from the Wrist Movement data (Figure 3) were trained on (7) wrist extension-flexion trials only or (8) wrist supination-pronation trials only (with pronation arbitrarily being analogous to flexion, i.e. having a kinematic value of 1, and supination therefore being analogous to extension, i.e. having a kinematic value of -1, in training).

To identify the alignment between those neural spaces, we then calculated the vector angle between the coefficients for every possible pair of models. Because we generated 5 extension-flexion regression models per category (5 folds of cross-validation), there were 25 vector angles per cross-categorical model pairing and 10 vector angles per within-category model pairing. The violin plots in Figure 4 represent the estimated probability density distribution of angles per categorical model pairing, the open dots depict the median, and the bars show the standard deviation. Colors are redundant and included only for ease of interpretation.

### Calculation of electrode modulation maps

We computed the mean peri-event time histogram in 20ms bins for each motor channel across a two second period centered on the start of movement. We then identified the response window during which the absolute difference between average 𝐹𝑅_𝑑𝑖𝑔𝑖𝑡_ (across repetitions) and 𝑚𝑒𝑎𝑛 (𝐹𝑅_𝑎𝑙𝑙_) (across repetitions) is at a maximum. Then,

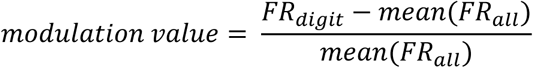

We repeated this process using 1s of the relaxation stage and computed the correlation between the activation map during movement and the activation map during relaxation. Channels where 𝑚𝑒𝑎𝑛 (𝐹𝑅_𝑎𝑙𝑙_) < 0.5 spikes/second were ignored and treated as NaNs.

### Modulation map significance

At each electrode on each array, we performed two Kruskal-Wallis tests on z-scored responses of the channel with digit identity and digit extension-flexion as factors (for Individual Finger Movement sessions) or one Kruskal-Wallis test with wrist orientation as the factor (for Wrist Movement sessions); the response data for each trial consists of the maximum z-scored response during the first 50% of the movement stage (when considering modulation significance during movement) or the maximum z-scored response during the first 50% of the relaxation stage (when considering modulation significance during relaxation). Spatial electrode modulation significance masks were created for movement during each session (Supp. Figure 1C, Supp. Figure 2C) by thresholding on the confidence (i.e. p-value) of the Kruskal-Wallis test for a given factor. Modulation was considered to be significant if p<0.05. Only significantly modulated electrodes were considered in the creation of Figure 1F and supplementary modulation maps. To assess the similarity of the positive modulation in array activity maps representative of individual fingers, the correlation between each pair of significance-masked maps (thumb vs. index, thumb vs. middle, etc.) was calculated for each session. These were compared against the distribution of correlations between trial-randomized maps within sessions. Wilcoxon rank sum tests were performed to establish the significance of the difference between distributions of correlations.

### Protocol for online control of individual fingers

At the start of each session, a regression decoder was trained on neural activity during 10 to 20 repeated observations of automated individual flexions and extensions of the thumb, index, and middle finger. During observation, the participant was instructed to observe and attempt or imagine performing the instructed movements. During online control, the participant was given 15s to move the audibly prompted target digit to its target position while holding all other fingers at the rest position (movement phase) and 15s to relax all digits to the neutral position once the target position was achieved (relaxation phase). To achieve a successful movement phase, the target digit had to remain within 30% of the target angle and each of the non-target digits had to remain within 30% of the neutral angle for 0.25s. To achieve a successful relaxation phase, the target digit and all non-target digits had to remain within 30% of the neutral angle for 0.25s. Trials proceeded to the relaxation phase only if the movement phase was successful. The hand avatar was automatically reset to the neutral position between each trial. To produce the trial-averaged traces in Figure 6A and Supp. Figure 7A, movement and relaxation phases were interpolated to the median duration of successful movement and relaxation phases across all trials.

### Protocol for online control of wrist movements

A second decoder, trained on neural activity during 10 to 20 repeated observations of automated wrist-only movements (neutral, flexion, extension, pronation, supination), was combined with the decoder previously derived from individual finger movement observations to produce a decoder capable of simultaneous prediction of individual digit position and wrist roll and pitch. During wrist-only online control, brain control of finger postures was deactivated, limiting the participant’s control space to wrist roll and pitch. As in digit-only online control, the participant was given 15s to assume the audibly prompted wrist orientation and 15s to relax to the neutral wrist position. To achieve successful movement or relaxation, wrist roll and pitch both had to be within 30% of their target values. The hand avatar was automatically reset to the neutral position between each trial. To produce the trial-averaged traces in Figure 6D and Supp. Figure 7C, movement and relaxation phases were interpolated to the median duration of successful movement and relaxation phases across all trials.

### Protocol for simultaneous online control of fingers and wrist

For simultaneous online control of fingers and wrist, brain control of both wrist roll and pitch and finger postures was enabled. During simultaneous online control, the participant was given 25s to move the target digit and wrist to the audibly prompted target positions while holding all other fingers at the rest position (movement phase) and 25s to relax all digits and the wrist to their respective neutral positions once the target positions were achieved (relaxation phase). As in online control of individual fingers, digit and wrist angles had to remain within 30% of their targets for 0.25s in order to achieve movement or relaxation phase success. To produce the trial-averaged traces in Supp. Figure 7HI, movement and relaxation phases were interpolated to the median duration of successful movement and relaxation phases across all trials.

### Online decoders

For online finger position decoding, a Kalman filter-based regression decoder was used. Principal component analysis (PCA) was performed on the 256 channels of neural activity recorded during observation and the first N components that could explain 90% of the variance in the data were used in training. The coefficient transformation matrix was stored for later application to incoming neural activity during online prediction. We extracted the angular position descriptive of each finger’s movement over the course of the observation period (ranging from -1 for full extension to +1 for full flexion) and combined those traces with the time courses of the first N PCs to produce a model of the common digit extension-flexion axis. After subtracting the common axis from neural activity, we trained a digit position-estimating Kalman filter. We also found the gradient of each of the five angular position traces to produce five corresponding angular velocity traces and combined those traces with the common axis-subtracted time courses of the first N PCs to produce a digit velocity-estimating Kalman filter. During online control, the estimated position of each digit was taken as the mean of the prediction from the position-estimating filter and the compounded position estimate from the velocity-estimating filter.

A similar procedure was repeated for online wrist control, focused on extracting the roll and pitch of the wrist using position- and velocity-estimating Kalman filters. The number and identities of the PCs explaining 90% of the variance in the data were reevaluated using the combined set of individual finger and wrist observation data. The common digit extension-flexion axis was recalculated in the new PC space and subtracted from neural activity prior to training and during online prediction.

### Subtracting the common extension-flexion axis from neural activity

The axis describing the common digit extension-flexion neural activity was calculated as

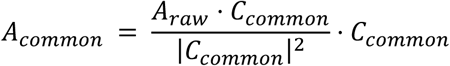

where A_raw_ is the PC-transformed incoming neural signal, 𝐶_𝑐𝑜𝑚𝑚𝑜𝑛_ are the estimated coefficients of the common digit extension-flexion axis, and · represents the dot product of vectors. In decoders with shared axis subtraction, the vector 𝐴_𝑐𝑜𝑚𝑚𝑜𝑛_ was subtracted from the PC-transformed incoming neural signal prior to Kalman filter training and prediction.

## Author contributions

AMXE, ARS, and SJB designed the study with input from other authors. AMXE, EVO, GHB, and ARS performed the experiments. AMXE and ARS performed the analysis. AMXE wrote the first draft of the manuscript. AMXE and ARS wrote the paper with input from all authors.

## Competing interests

The authors declare no competing interests.

## Acknowledgements

The authors would like to thank the participants for their time. The authors would like to thank Sensorimotor Bionics Group at the University of Chicago and interuniversity Cortical Bionics Research Group for insightful discussions and technical support. This work was supported by the National Institute of Neurological Disorders and Stroke, R35 NS122333, R01 NS131953 and R01 NS130302.

## Data availability

All data generated or analyzed during this study as well as analysis and plotting scripts are going to be made available on DABI upon publication.

## Code availability

All analysis and plotting scripts are going to be made available in a public repository https://github.com/SobinovLab/finger_individuation

## Supplementary Materials

**Supp. Figure 1.**
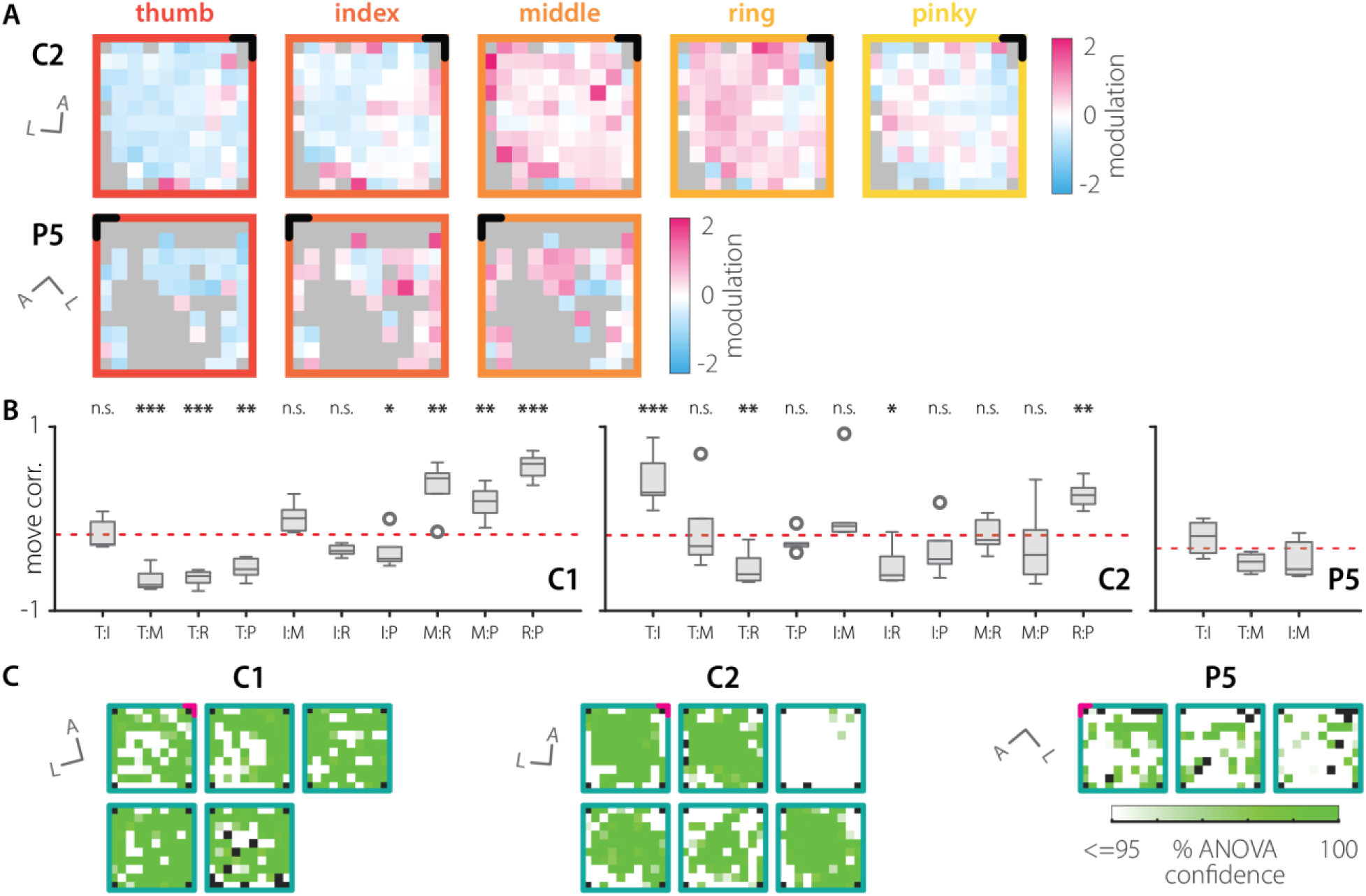
**A|** Modulation by digit identity (thumb, index, middle, ring, and pinky, red to yellow borders) in C2’s lateral motor array during solitary digit movement, averaged across sessions. Modulation by digit identity in P5’s medial motor array during solitary digit movement, averaged across sessions. Non-significant channels are gray. Anterior-most medial corner of arrays is black. **B|** Distributions (across sessions) of correlation of the modulation of array activity maps of the annotated pair of digits (T, thumb; I, index; M, middle; R, ring; P, pinky) in C1, C2, and P5. p-values represent the result of a significance test between the correlation distribution for trial-randomized maps (mean per subject presented as red dotted line; C1: -0.17, C2: -0.18, P5: -0.33) and the denoted pair of maps (R:P, ring vs. pinky). Top and bottom edges of each box are 75^th^ and 25^th^ percentiles. Line mid-box is median. Whiskers present maximum and minimum non-outliers. Outliers are represented as separate dots. * p<0.05, ** p<0.01, *** p<0.001. (C1 n=5, C2 n=6, P5 n=3) **C|** Kruskal-Wallis confidence in significance of electrode modulation by digit identity during movement in the lateral array (C1, C2) or medial array (P5) during individual finger movement sessions. p = (100-confidence)/100. See Methods for details. Anterior-most medial corner of arrays is pink.

**Supp. Figure 2.**
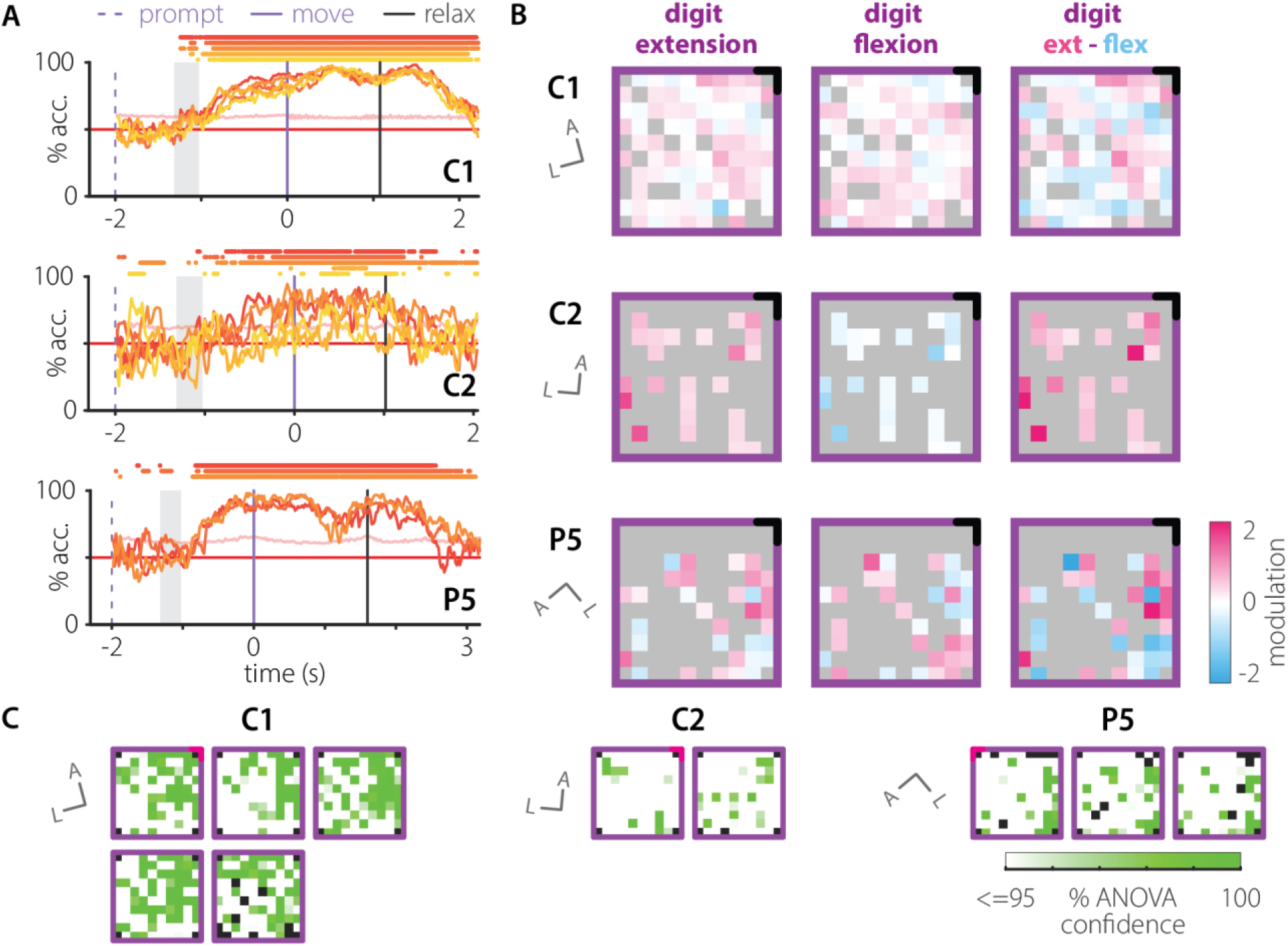
**A|** Classification of digit movement direction – flexion or extension – for individual digits. Colored bars above traces correspond to points significantly deviating from chance (red line). Pale pink line depicts chance significance threshold. (C1 n=5, C2 n=2, P5 n=3). **B|** Digit movement direction modulation array activity maps for digit extension (left), digit flexion (center), and digit extension minus flexion (right) in C1, C2, and P5. Non-significant channels are gray. Anterior-most medial corner of arrays is black. **C|** Kruskal-Wallis confidence in significance of electrode modulation by digit extension-flexion during movement in the lateral array (C1, C2) or medial array (P5) during individual finger movement sessions. p = (100-confidence)/100. See Methods for details. Anterior-most medial corner of arrays is pink.

**Supp. Figure 3.**
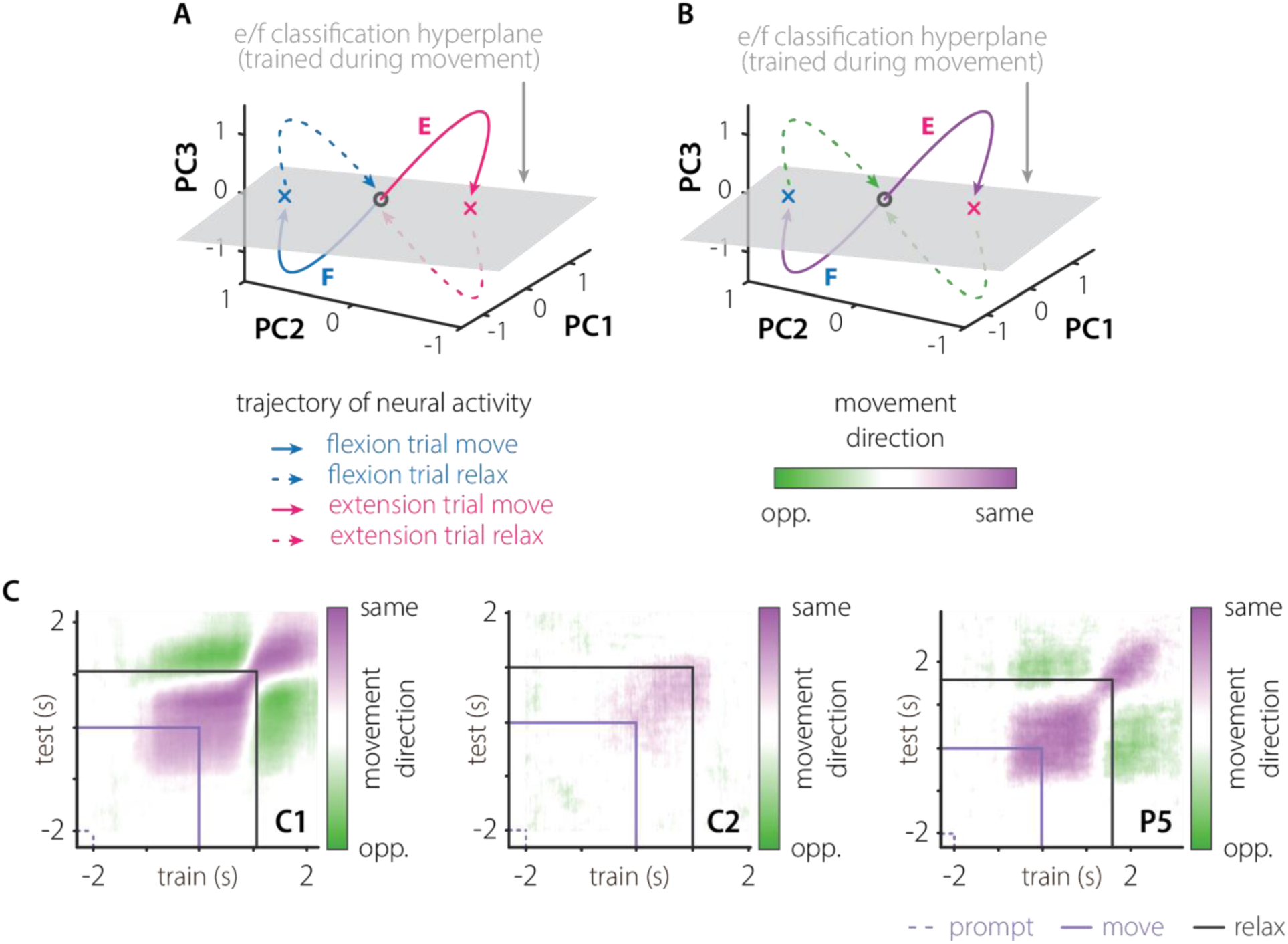
**A|** The placement of the linear classifier’s extension-flexion hyperplane in neural space is such that the flexion during relaxation in an extension trial (pink dotted line) is more similar to the flexion during movement in a flexion trial (blue solid line) than it is to the extension during movement in an extension trial (pink solid line). **B|** Consequently, movement occurring during the relaxation phase is reliably classified as opposite the trial type (green coloration on dotted lines). **C|** Temporal generalization heatmaps of digit movement direction for C1, C2, and P5, averaged across digits. Color corresponds to the classification of the instructed movement direction (purple) or movement in the opposite direction (green). In C1 and P5, any single finger flexion or extension is a composite movement of both flexion and extension.

**Supp. Figure 4.**
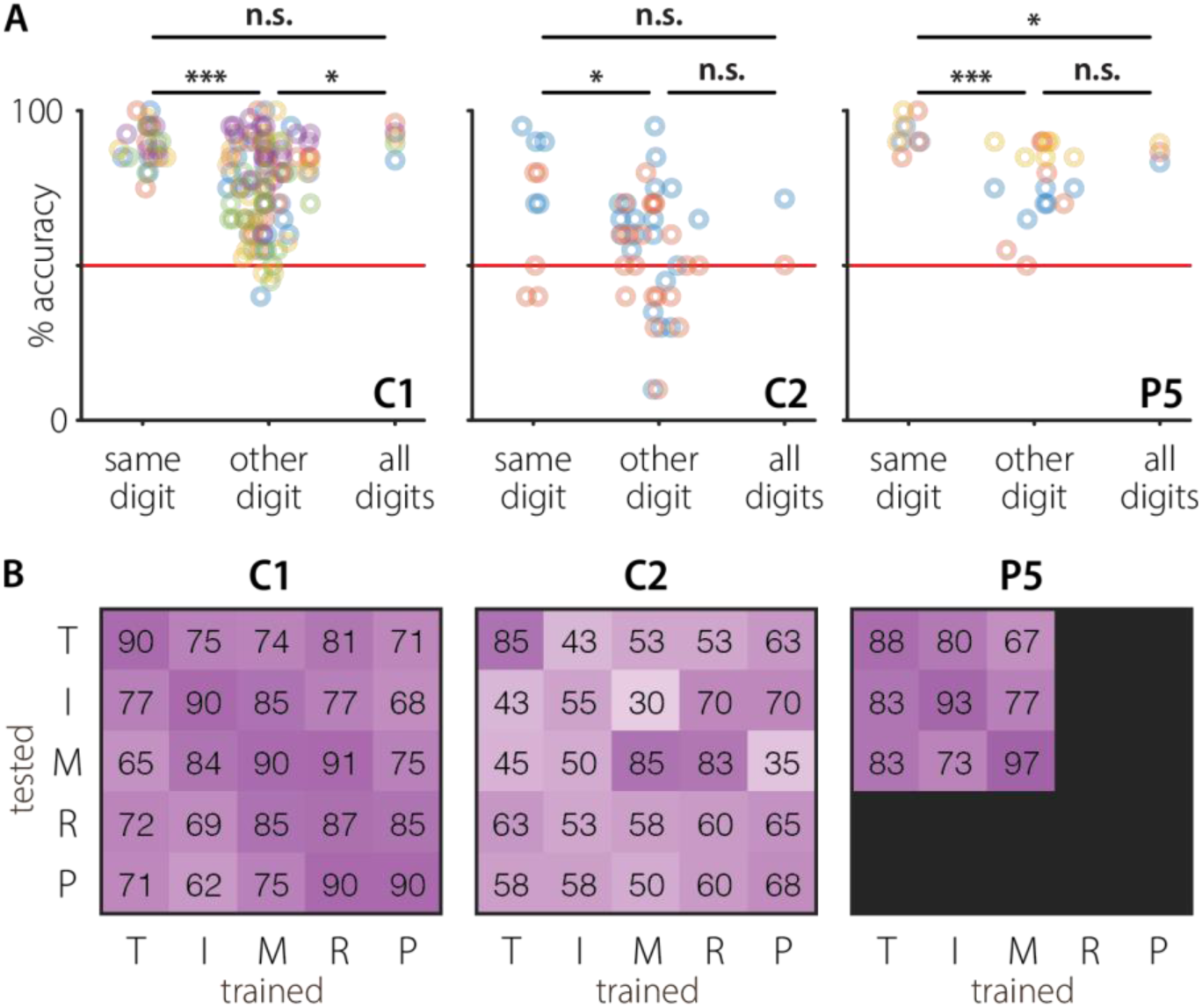
**A|** Generalization of movement direction across digits during movement phase in subjects C1, C2, and P5. The extension-flexion classifier was trained on a single digit and tested on the same digit (“same digit”) or a different digit (“other digit”) or trained on a pooled dataset containing all digits and tested on held-out examples of all digits (“all digits”). Each dot represents accuracy in a single session for a specific training and testing digit combination. Dot color corresponds to session. Horizontal solid red line illustrates chance level. * p<0.05, ** p<0.01, *** p<0.001. **B|** Average digit extension-flexion classification accuracy across sessions for all combinations of training and testing digit identity in C1, C2, and P5. A strong diagonal suggests that extension-flexion decoding is most successful when training and testing on the same digit, but classification accuracies well above chance (50%) for other train/test digit pairs provide evidence for a shared extension-flexion classification axis across digits.

**Supp. Figure 5.**
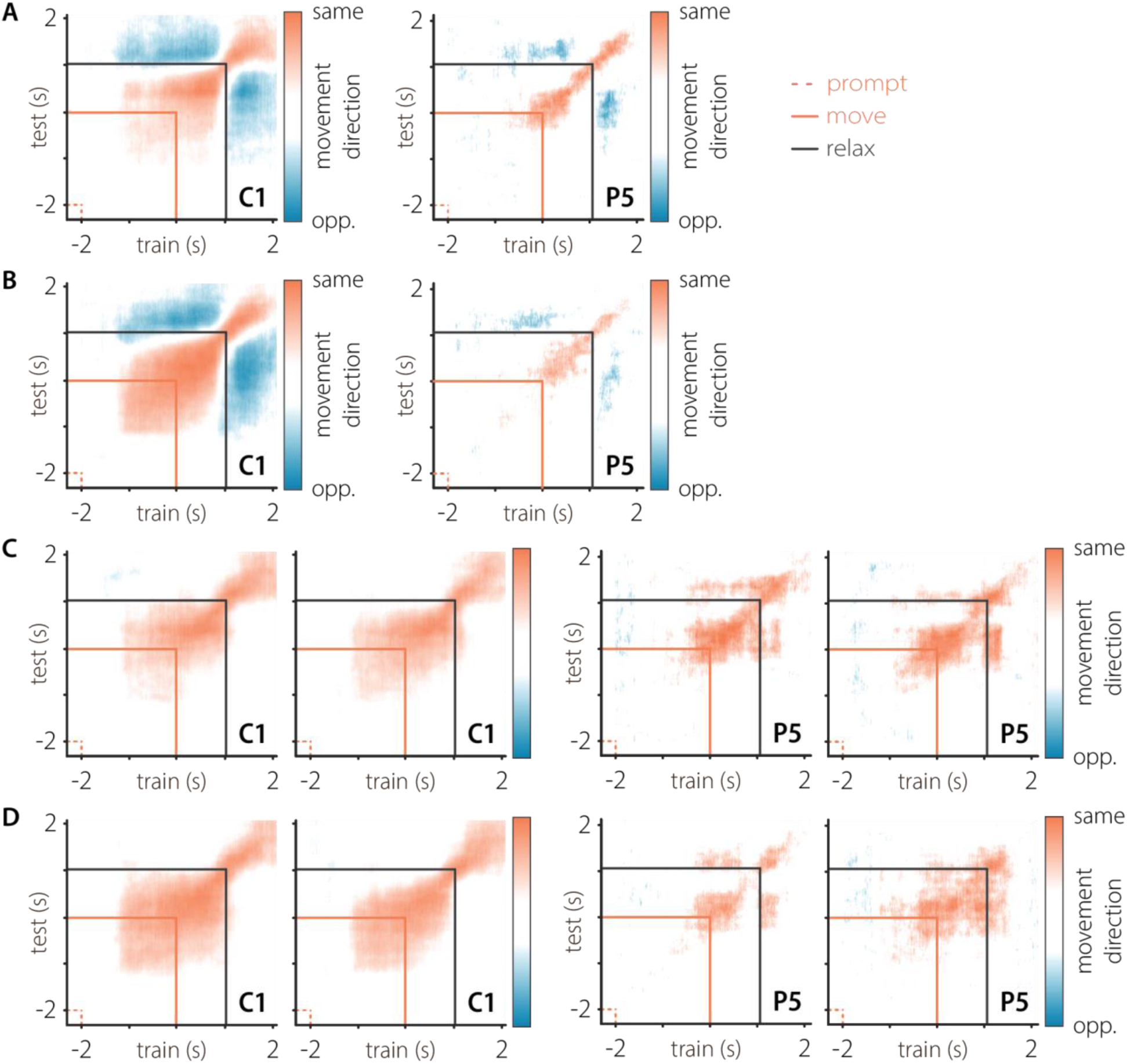
**A|** Temporal generalization heatmap of wrist extension-flexion movement direction for C1 and P5. Color corresponds to instructed (orange) or opposite (blue) direction classification. A wrist orientation classifier trained on the movement stage or the relaxation stage correctly identifies the motion occurring during relaxation as the inverse of the motion occurring during movement. **B|** Same as A for supination-pronation axis. **C|** Distinct phases in A are not visible in C1 or P5 when performing analysis using non-complementary wrist orientations (left in subject pair: neutral-extension, right in subject pair: neutral-flexion). **D|** Distinct phases in B are not visible in C1 or P5 when performing analysis using non-complementary wrist orientations (left in subject pair: neutral-pronation, right in subject pair: neutral-supination).

**Supp. Figure 6.**
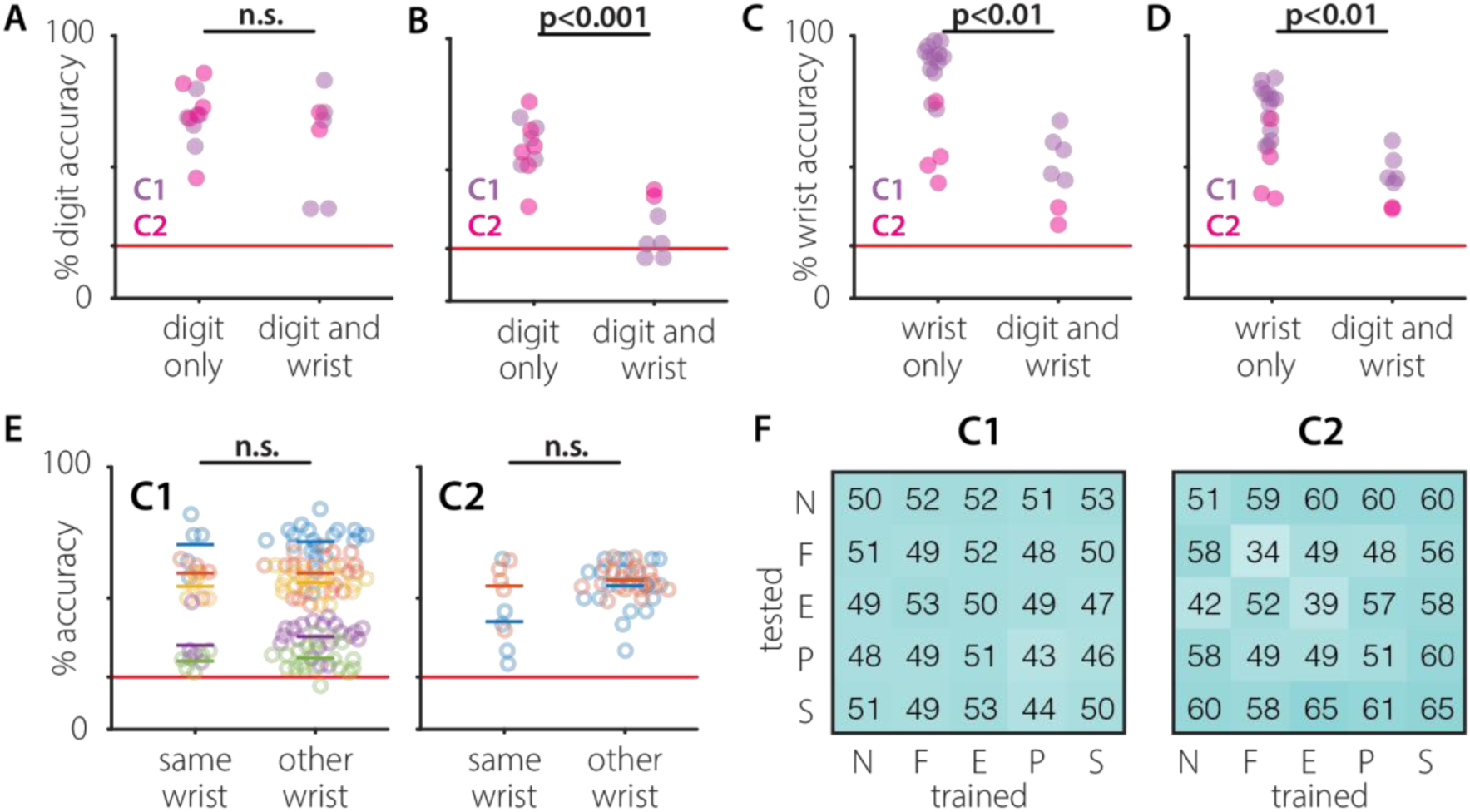
**A|** Distribution of maximum accuracy of digit classification during movement in digit-only trials and simultaneous digit and wrist trials (C1: purple, C2: pink). **B|** Distribution of maximum accuracy of digit classification during the preparatory period in digit-only trials and simultaneous digit and wrist trials. Colors as in A. **C|** Same as A for wrist orientation classification. **D|** Same as B for wrist orientation classification. **E|** Digit classification training and testing at different wrist orientations, color-coded by session. The wide span of digit classification accuracies is due to large variations in overall performance across sessions, not due to deficits specific to training/testing wrist combinations within a session. Consistent and mixed wrist swarms are not significantly different within any single session. (C1 n=5, C2 n=2) **F|** Accuracy of digit classification in C1 when trained within one wrist orientation and tested on another for C1 and C2.

**Supp. Figure 7.**
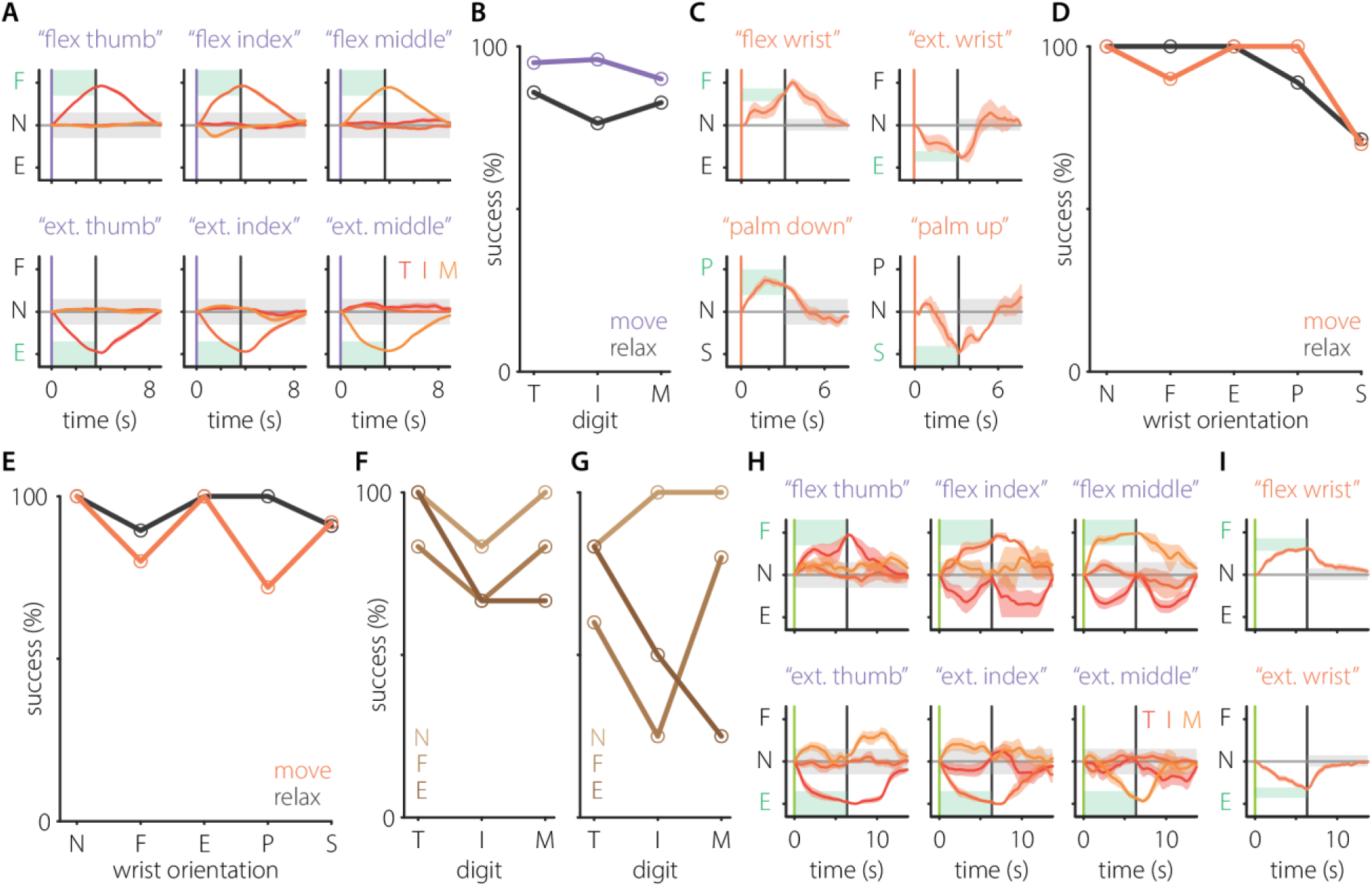
**A|** Position during successful individual finger movement trials completed using a decoder without shared axis subtraction, averaged by target (C1 n=4) and interpolated to the median stage length across successful trials. Target window is shaded green. Gray regions represent error threshold for non-target digits and for target digit during return. Red and orange shaded regions surrounding traces depict the standard error of the mean. **B|** Trial success rates during movement (purple trace) and relaxation (gray trace) phases for trials in A, broken down by digit. **C|** Extension/flexion (top row) and pronation/supination (bottom row) during successful wrist movement trials completed using a decoder without shared axis subtraction, averaged by target (C1 n=1). Interpolation and target window shading as in A. Orange regions surrounding traces depict the standard error of the mean. Median time to movement success 2.3s (IQR 0.9-3.3s). Median time to relaxation success 2.8s (IQR 1.4-4.4s). **D|** Same as B, but for wrist control trials in C. Move success 90% (36/40). Relax success 91% (31/34). **E|** Same as D, but for wrist control trials completed using a decoder with shared axis subtraction. **F|** Digit success rates during the movement phase of simultaneous wrist and finger movements, grouped by wrist motion and broken down by digit. **G|** F, for the relaxation phase of simultaneous wrist and finger movements. **H|** Digit position during successful simultaneous wrist and finger movement trials completed using a decoder with shared axis subtraction, averaged by target (C1 n=1) and interpolated to the median stage length across successful trials. Colors as in A. **I|** Wrist movements during successful simultaneous wrist and finger trials, with brain control of wrist at 20%.

## Notes

### Competing Interest Statement

The authors have declared no competing interest.

### Summary of Updates

Fix to the figure numbering in Supplements

## References

1. Acharya, S. et al. (2007) “Towards a Brain-Computer Interface for Dexterous Control of a Multi-Fingered Prosthetic Hand,” 2007 3rd International IEEE/EMBS Conference on Neural Engineering. 2007 3rd International IEEE/EMBS Conference on Neural Engineering, Kohala Coast, HI, USA: IEEE, pp. 200–203. Available at: 10.1109/CNE.2007.369646.

2. Adewuyi, A.A., Hargrove, L.J. and Kuiken, T.A. (2017) “Resolving the effect of wrist position on myoelectric pattern recognition control,” Journal of NeuroEngineering and Rehabilitation, 14(1), p. 39. Available at: 10.1186/s12984-017-0246-x.

3. Aggarwal, V. et al. (2008) “Asynchronous Decoding of Dexterous Finger Movements Using M1 Neurons,” IEEE Transactions on Neural Systems and Rehabilitation Engineering, 16(1), pp. 3–14. Available at: 10.1109/TNSRE.2007.916289.

4. Aggarwal, V. et al. (2013) “State-based decoding of hand and finger kinematics using neuronal ensemble and LFP activity during dexterous reach-to-grasp movements,” Journal of Neurophysiology, 109(12), pp. 3067–3081. Available at: 10.1152/jn.01038.2011.

5. Agudelo-Toro, A. et al. (2024) “Accurate neural control of a hand prosthesis by posture-related activity in the primate grasping circuit,” Neuron, 0(0). Available at: 10.1016/j.neuron.2024.09.018.

6. Ajiboye, A.B. et al. (2012) “Prediction of Imagined Single-Joint Movements in a Person with High Level Tetraplegia,” IEEE transactions on bio-medical engineering, 59(10), pp. 2755–2765. Available at: 10.1109/TBME.2012.2209882.

7. Ajiboye, A.B. et al. (2017) “Restoration of reaching and grasping movements through brain-controlled muscle stimulation in a person with tetraplegia: a proof-of-concept demonstration.,” *Lancet (London*, England), 389(10081), pp. 1821–1830. Available at: 10.1016/S0140-6736(17)30601-3.

8. Anderson, K.D. (2004) “Targeting Recovery: Priorities of the Spinal Cord-Injured Population,” Journal of Neurotrauma, 21(10), pp. 1371–1383. Available at: 10.1089/neu.2004.21.1371.

9. Baldwin, M.K.L. et al. (2018) “Representations of fine digit movements in posterior and anterior parietal cortex revealed using long-train intracortical microstimulation in macaque monkeys,” Cerebral Cortex, 28(12), pp. 4244–4263. Available at: 10.1093/cercor/bhx279.

10. Beringer, C.R. et al. (2020) “The effect of wrist posture on extrinsic finger muscle activity during single joint movements,” Scientific Reports, 10(1), p. 8377. Available at: 10.1038/s41598-020-65167-x.

11. Branco, M.P. et al. (2019) “Encoding of kinetic and kinematic movement parameters in the sensorimotor cortex: A Brain-Computer Interface perspective,” European Journal of Neuroscience, 50(5), pp. 2755–2772. Available at: 10.1111/ejn.14342.

12. Capaday, C. et al. (2013) “On the functional organization and operational principles of the motor cortex,” Frontiers in Neural Circuits, 7. Available at: 10.3389/fncir.2013.00066.

13. Carandini, M. and Heeger, D.J. (2012) “Normalization as a canonical neural computation,” Nature Reviews Neuroscience, 13(1), pp. 51–62. Available at: 10.1038/nrn3136.

14. Chehade, N.G. and Gharbawie, O.A. (2023) “Motor actions are spatially organized in motor and dorsal premotor cortex,” *eLife*. Edited by J. Diedrichsen, J.I. Gold, and W. Vanduffel, 12, p. e83196. Available at: 10.7554/eLife.83196.

15. Collinger, J.L., Boninger, M.L., et al. (2013) “Functional priorities, assistive technology, and brain-computer interfaces after spinal cord injury,” The Journal of Rehabilitation Research and Development, 50(2), p. 145. Available at: 10.1682/JRRD.2011.11.0213.

16. Collinger, J.L., Wodlinger, B., et al. (2013) “High-performance neuroprosthetic control by an individual with tetraplegia,” The Lancet, 381(9866), pp. 557–564. Available at: 10.1016/S0140-6736(12)61816-9.

17. Dechent, P. and Frahm, J. (2003) “Functional somatotopy of finger representations in human primary motor cortex,” Human Brain Mapping, 18(4), pp. 272–283. Available at: 10.1002/hbm.10084.

18. Dekleva, B.M. and Collinger, J.L. (2025) “Using transient, effector-specific neural responses to gate decoding for brain–computer interfaces,” Journal of Neural Engineering, 22(1), p. 016036. Available at: 10.1088/1741-2552/adaa1f.

19. Donoghue, J.P., Leibovic, S. and Sanes, J.N. (1992) “Organization of the forelimb area in squirrel monkey motor cortex: representation of digit, wrist, and elbow muscles,” Experimental Brain Research, 89(1). Available at: 10.1007/BF00228996.

20. Duinen, H. van and Gandevia, S.C. (2011) “Constraints for control of the human hand,” The Journal of Physiology, 589(23), pp. 5583–5593. Available at: 10.1113/jphysiol.2011.217810.

21. Ejaz, N., Hamada, M. and Diedrichsen, J. (2015) “Hand use predicts the structure of representations in sensorimotor cortex.,” Nature neuroscience, 18(7), pp. 1034–40. Available at: 10.1038/nn.4038.

22. Ferguson, K.A. and Cardin, J.A. (2020) “Mechanisms underlying gain modulation in the cortex,” Nature reviews. Neuroscience, 21(2), pp. 80–92. Available at: 10.1038/s41583-019-0253-y.

23. Flesher, S.N. et al. (2021) “A brain-computer interface that evokes tactile sensations improves robotic arm control,” Science, 372(6544), pp. 831–836. Available at: 10.1126/science.abd0380.

24. Georgopoulos, A.P. et al. (1999) “Neural coding of finger and wrist movements,” Journal of Computational Neuroscience, 6(3), pp. 279–288. Available at: 10.1023/a:1008810007672.

25. Goodman, J.M. et al. (2019) “Postural Representations of the Hand in the Primate Sensorimotor Cortex.,” Neuron, p. 566539. Available at: 10.1016/j.neuron.2019.09.004.

26. Guan, C. et al. (2023) “Decoding and geometry of ten finger movements in human posterior parietal cortex and motor cortex,” Journal of Neural Engineering, 20(3), p. 036020. Available at: 10.1088/1741-2552/acd3b1.

27. Hager-Ross, C. and Schieber, M.H. (2000) “Quantifying the independence of human finger movements: Comparisons of digits, hands, and movement frequencies,” Journal of Neuroscience, 20(22), pp. 8542–8550. Available at: 10.1523/jneurosci.20-22-08542.2000.

28. Hluštík, P. et al. (2001) “Somatotopy in Human Primary Motor and Somatosensory Hand Representations Revisited,” Cerebral Cortex, 11(4), pp. 312–321. Available at: 10.1093/cercor/11.4.312.

29. Hotson, G., McMullen, D.P., Fifer, Matthew S, et al. (2016) “Individual finger control of a modular prosthetic limb using high-density electrocorticography in a human subject,” Journal of Neural Engineering, 13(2), p. 026017. Available at: 10.1088/1741-2560/13/2/026017.

30. Hotson, G., McMullen, D.P., Fifer, Matthew S., et al. (2016) “Individual Finger Control of the Modular Prosthetic Limb using High-Density Electrocorticography in a Human Subject,” Journal of neural engineering, 13(2), p. 026017. Available at: 10.1088/1741-2560/13/2/026017.

31. Indovina, I. and Sanes, J.N. (2001) “On Somatotopic Representation Centers for Finger Movements in Human Primary Motor Cortex and Supplementary Motor Area,” NeuroImage, 13(6), pp. 1027–1034. Available at: 10.1006/nimg.2001.0776.

32. Irwin, Z.T. et al. (2017) “Neural control of finger movement via intracortical brain–machine interface,” Journal of Neural Engineering, 14(6), p. 066004. Available at: 10.1088/1741-2552/aa80bd.

33. Jorge, A. et al. (2020) “Classification of Individual Finger Movements Using Intracortical Recordings in Human Motor Cortex,” Neurosurgery, 87(4), pp. 630–638. Available at: 10.1093/neuros/nyaa026.

34. Jude, J.J. et al. (2026) “Restoring rapid natural bimanual typing with a neuroprosthesis after paralysis,” Nature Neuroscience, pp. 1–10. Available at: 10.1038/s41593-026-02218-y.

35. Kilbreath, S.L. and Gandevia, S.C. (1994) “Limited independent flexion of the thumb and fingers in human subjects.,” The Journal of Physiology, 479(3), pp. 487–497. Available at: 10.1113/jphysiol.1994.sp020312.

36. Kirsch, E., Rivlis, G. and Schieber, M.H. (2014) “Primary Motor Cortex Neurons during Individuated Finger and Wrist Movements: Correlation of Spike Firing Rates with the Motion of Individual Digits versus Their Principal Components,” Frontiers in Neurology, 5, p. 70. Available at: 10.3389/fneur.2014.00070.

37. Kubánek, J. et al. (2009) “Decoding flexion of individual fingers using electrocorticographic signals in humans,” Journal of Neural Engineering, 6(6), p. 066001. Available at: 10.1088/1741-2560/6/6/066001.

38. Lang, C.E. and Schieber, M.H. (2004) “Reduced Muscle Selectivity During Individuated Finger Movements in Humans After Damage to the Motor Cortex or Corticospinal Tract,” Journal of Neurophysiology, 91(4), pp. 1722–1733. Available at: 10.1152/jn.00805.2003.

39. Lemelin, P. and Diogo, R. (2016) “Anatomy, Function, and Evolution of the Primate Hand Musculature,” in T.L. Kivell et al. (eds.) The Evolution of the Primate Hand. New York.

40. Lemon, R.N. and Griffiths, J. (2005) “Comparing the function of the corticospinal system in different species: Organizational differences for motor specialization?,” Muscle and Nerve, 32(3), pp. 261–279. Available at: 10.1002/mus.20333.

41. Mender, M.J. et al. (2024) “Functional Electrical Stimulation and Brain-Machine Interfaces for Simultaneous Control of Wrist and Finger Flexion.” Neuroscience. Available at: 10.1101/2024.08.11.607263.

42. Miller, K.J. et al. (2009) “Decoupling the Cortical Power Spectrum Reveals Real-Time Representation of Individual Finger Movements in Humans,” Journal of Neuroscience, 29(10), pp. 3132–3137. Available at: 10.1523/JNEUROSCI.5506-08.2009.

43. Mohamed, A.-K. and Aharonson, V. (2021) “Four-class BCI discrimination of right and left wrist and finger movements,” IFAC-PapersOnLine, 54(21), pp. 91–96. Available at: 10.1016/j.ifacol.2021.12.016.

44. Moran, D.W. and Schwartz, A.B. (1999) “Motor cortical representation of speed and direction during reaching,” Journal of Neurophysiology, 82, pp. 2676–2692.

45. Nason, S.R. et al. (2021) “Real-time linear prediction of simultaneous and independent movements of two finger groups using an intracortical brain-machine interface,” Neuron, 109(19), pp. 3164–3177.e8. Available at: 10.1016/j.neuron.2021.08.009.

46. Nason-Tomaszewski, S.R. et al. (2023) “Restoring continuous finger function with temporarily paralyzed nonhuman primates using brain–machine interfaces,” Journal of Neural Engineering, 20(3), p. 036006. Available at: 10.1088/1741-2552/accf36.

47. Okorokova, E.V. et al. (2020) “Decoding hand kinematics from population responses in sensorimotor cortex during grasping,” Journal of Neural Engineering, 17(4), p. 046035. Available at: 10.1088/1741-2552/ab95ea.

48. Paninski, L. et al. (2004) “Spatiotemporal Tuning of Motor Cortical Neurons for Hand Position and Velocity,” Journal of Neurophysiology [Preprint]. Available at: 10.1152/jn.00587.2002.

49. Penfield, W. and Rasmussen, T. (1950) The cerebral cortex of man: a clinical study of localization of function. New York: Macmillan.

50. Popp, W.L. et al. (2023) “Effects of wrist posture and stabilization on precision grip force production and muscle activation patterns,” Journal of Neurophysiology [Preprint]. Available at: 10.1152/jn.00420.2020.

51. Rasmussen, R.G., Schwartz, A. and Chase, S.M. (2017) “Dynamic range adaptation in primary motor cortical populations,” eLife, 6, p. e21409. Available at: 10.7554/eLife.21409.

52. Rastogi, A. et al. (2021) “The Neural Representation of Force across Grasp Types in Motor Cortex of Humans with Tetraplegia,” eNeuro, 8(1). Available at: 10.1523/ENEURO.0231-20.2020.

53. Rathelot, J.A. and Strick, P.L. (2006) “Muscle representation in the macaque motor cortex: An anatomical perspective,” Proceedings of the National Academy of Sciences of the United States of America, 103(21), pp. 8257–8262. Available at: 10.1073/pnas.0602933103.

54. Rathelot, J.A. and Strick, P.L. (2009) “Subdivisions of primary motor cortex based on cortico-motoneuronal cells,” Proceedings of the National Academy of Sciences of the United States of America, 106(3), pp. 918–923. Available at: 10.1073/pnas.0808362106.

55. Reilly, K.T. and Schieber, M.H. (2003) “Incomplete functional subdivision of the human multitendoned finger muscle flexor digitorum profundus: An electromyographic study,” Journal of Neurophysiology, 90(4), pp. 2560–2570. Available at: 10.1152/jn.00287.2003.

56. Rizzolatti, G. and Luppino, G. (2001) “The Cortical Motor System,” Neuron, 31(6), pp. 889–901. Available at: 10.1016/S0896-6273(01)00423-8.

57. Schieber, M.H. (2001) “Constraints on somatotopic organization in the primary motor cortex,” Journal of Neurophysiology, 86(5), pp. 2125–2143. Available at: 10.1152/jn.2001.86.5.2125.

58. Schieber, M.H. and Hibbard, L.S. (1993) “How somatotopic is the motor cortex hand area?,” Science, 261(5120), pp. 489–492. Available at: 10.1126/science.8332915.

59. Schieber, M.H. and Santello, M. (2004) “Hand function: Peripheral and central constraints on performance,” Journal of Applied Physiology, 96(6), pp. 2293–2300. Available at: 10.1152/japplphysiol.01063.2003.

60. Seo, N.J. et al. (2008) “Wrist strength is dependent on simultaneous power grip intensity.,” Ergonomics, 51(10), pp. 1594–1605. Available at: 10.1080/00140130802216925.

61. Shah, N.P. et al. (2023) “A brain-computer typing interface using finger movements,” 2023 11th International IEEE/EMBS Conference on Neural Engineering (NER). 2023 11th International IEEE/EMBS Conference on Neural Engineering (NER), Baltimore, MD, USA: IEEE, pp. 1–4. Available at: 10.1109/NER52421.2023.10123912.

62. Shelchkova, N.D. et al. (2023) “Microstimulation of human somatosensory cortex evokes task-dependent, spatially patterned responses in motor cortex,” Nature Communications, 14(1), p. 7270. Available at: 10.1038/s41467-023-43140-2.

63. Sobinov, A.R. and Bensmaia, S.J. (2021) “The neural mechanisms of manual dexterity,” Nature Reviews Neuroscience, pp. 1–17. Available at: 10.1038/s41583-021-00528-7.

64. Takei, T. et al. (2017) “Neural basis for hand muscle synergies in the primate spinal cord,” Proceedings of the National Academy of Sciences, 114(32), pp. 8643–8648. Available at: 10.1073/pnas.1704328114.

65. Takei, T. and Seki, K. (2010) “Spinal Interneurons Facilitate Coactivation of Hand Muscles during a Precision Grip Task in Monkeys,” Journal of Neuroscience, 30(50), pp. 17041–17050. Available at: 10.1523/JNEUROSCI.4297-10.2010.

66. Vargas-Irwin, C.E. et al. (2010) “Decoding Complete Reach and Grasp Actions from Local Primary Motor Cortex Populations,” Journal of Neuroscience, 30(29), pp. 9659–9669. Available at: 10.1523/JNEUROSCI.5443-09.2010.

67. Vargas-Irwin, C.E. et al. (2024) “Gesture encoding in human left precentral gyrus neuronal ensembles.” Neuroscience. Available at: 10.1101/2024.08.23.608325.

68. Vaskov, A.K. et al. (2018) “Cortical Decoding of Individual Finger Group Motions Using ReFIT Kalman Filter,” Frontiers in Neuroscience, 12. Available at: https://www.frontiersin.org/articles/10.3389/fnins.2018.00751 (Accessed: August 11, 2023).

69. Wang, W. et al. (2007) “Motor Cortical Representation of Position and Velocity During Reaching,” Journal of Neurophysiology, 97(6), pp. 4258–4270. Available at: 10.1152/jn.01180.2006.

70. Wang, W. et al. (2009) “Human Motor Cortical Activity Recorded with Micro-ECoG Electrodes During Individual Finger Movements,” Conference proceedings : … Annual International Conference of the IEEE Engineering in Medicine and Biology Society. IEEE Engineering in Medicine and Biology Society. Conference, 2009, pp. 586–589. Available at: 10.1109/IEMBS.2009.5333704.

71. Weber, D.J. et al. (2011) “Limb-state information encoded by peripheral and central somatosensory neurons: Implications for an afferent interface,” IEEE Transactions on Neural Systems and Rehabilitation Engineering, 19(5), pp. 501–513. Available at: 10.1109/TNSRE.2011.2163145.

72. Willsey, M.S. et al. (2025) “A high-performance brain–computer interface for finger decoding and quadcopter game control in an individual with paralysis,” Nature Medicine, 31(1), pp. 96–104. Available at: 10.1038/s41591-024-03341-8.

73. Wodlinger, B. et al. (2015) “Ten-dimensional anthropomorphic arm control in a human brain-machine interface: difficulties, solutions, and limitations.,” Journal of neural engineering, 12(1), p. 016011. Available at: 10.1088/1741-2560/12/1/016011.

74. Yan, Y., Sobinov, A.R. and Bensmaia, S.J. (2022) “Prehension kinematics in humans and macaques,” Journal of Neurophysiology [Preprint]. Available at: 10.1152/jn.00522.2021.

75. Zatsiorsky, V.M., Li, Z.-M. and Latash, M.L. (2000) “Enslaving effects in multi-finger force production,” Experimental Brain Research, 131(2), pp. 187–195. Available at: 10.1007/s002219900261.

